# Genomic Insights into the Demographic History and Local Adaptation of Wild Boars Across Eurasia

**DOI:** 10.1101/2025.02.05.636574

**Authors:** Zishuai Wang, Zixin Li, Tao Huang, Jianhai Chen, Pan Xu, Ruimin Qiao, Hongwei Yin, Chengyi Song, Dongjie Zhang, Di Liu, Shuhong Zhao, Martien A. M. Groenen, Ole Madsen, Yanlin Zhang, Lijing Bai, Kui Li

## Abstract

Wild boars exhibit genetic and phenotypic diversity shaped by migrations and local adaptations. Its expansion across Eurasia, especially in central Asia, remains underexplored. Here, we present newly sequenced whole-genome data of 47 wild boars from Eastern Asia, Central Asia, and Europe, combined with 49 existing genomes, creating a comprehensive dataset of 96 individuals. Our analyses show that Asian wild boars and Southeast Asian Suids split ∼3.6 Mya, with Central Asian and Southern Chinese ancestors diverging ∼1.8 Mya. The split between Central Asian and European-Near East ancestors occurred ∼0.9 Mya, followed by a European-Near East divergence ∼0.6 Mya. We identify signatures of local adaptation in Central Asian populations, including two positively selected variants in *LPIN1*, associated with cold adaptation, and a missense mutation in *ALPK2*, linked to meat traits. These findings provide insights into wild boar dispersal and adaptation, and shed light on domestic pigs breeding.

## Introduction

*Sus scrofa*, commonly known as the wild boar^1^, is an ecologically and agriculturally important species with a significant impact on ecosystems and human economies^2^. Emerging in Southeast Asia around 3 to 4 million years ago^3, 4, 5^, *S. scrofa* has expanded its range across Eurasia and into North Africa, demonstrating remarkable adaptability to diverse environmental conditions^6, 7, 8, 9, 10^. Physically robust, with powerful limbs and sharp tusks, *S. scrofa* occupies a wide range of habitats, from dense forests to open grasslands. Its omnivorous and opportunistic diet, which includes roots, fruits, small animals, and carrion, enables it to exploit various food resources and adapt to seasonal fluctuations. Socially, wild boars form cohesive family groups that support cooperative foraging and predator defense. Environmental factors limiting its range expansion include deep snow in winter^7^ and the arid climates of the Gobi Desert and Mongolian steppe^11^. This wide geographic distribution has resulted in extensive genetic diversity and ecological flexibility, making the species a valuable model for studying population genetics, evolutionary history and adaptation.

Southeast Asia, particularly Island Southeast Asia (ISEA) and mainland Southeast Asia (MSEA), is considered the center of origin for *S. scrofa* and a biodiversity hotspot for the genus *Sus*^12, 13^. Wild boars from MSEA, especially the “Mekong region” identified by Wu et al.^14^, show high genetic diversity and encompass nearly all major East Asian mitochondrial haplotypes. Molecular phylogenetic studies have consistently indicated a significant divergence between Asian and European wild boar populations, estimated to have occurred during the Mid-Pleistocene, approximately 1.6 to 0.8 million years ago^15, 16, 17, 18, 19^. Despite considerable progress in characterizing the genetic structure of wild boars across East Asia^14, 20, 21^, little is known about the populations in Central Asia. Central Asian wild boars, situated at the crossroads of East Asia and Europe, likely played a role in gene flow and migration between distinct wild boar lineages, providing key insights into the dispersal patterns that contributed to the spread of *Sus scrofa* across these regions. Based on mitochondrial data, a recent study proposed migration routes of *Sus scrofa* populations from Asia to Europe^22^. However, whole-genome analyses are needed to resolve the remaining uncertainties. This gap in knowledge underscores the need for further research to clarify the dispersal patterns and genetic diversity that have shaped the *Sus scrofa* populations across this vast region.

In this study, we aim to clarify the dispersal history and genetic diversity of *Sus scrofa* across its broad geographic range, focusing on underrepresented Central Asian populations. Using whole-genome data integrated with existing datasets, we investigate the evolutionary relationships and migration routes that have shaped wild boar populations from Southeast Asia to Europe. Our analysis further seeks to identify genomic regions associated with local adaptation in Central Asia, providing insights into the role of natural selection in shaping the genetic landscape of these populations. These findings enhance our understanding of the evolutionary dynamics and adaptive processes of Sus scrofa across diverse ecological and geographic contexts.

## Results

### Sample collection and Genome-wide variation in wild boars

We sequenced the genomes of 47 wild boars across Eastern Asia, Central Asia—a region previously underrepresented in genetic studies—and Europe, achieving an average genomic coverage exceeding 30-fold. To establish a comprehensive view of genetic diversity among wild boars, these new data were integrated with 51 publicly available genomes (Supplementary Fig. S1, Supplementary Table 1). Additionally, six whole-genome sequences from another wild *Sus* species (*Sus cebifrons*) were included to facilitate demographic history analyses. This dataset encompasses a wide geographic range, representing 52 wild boars from Eastern Asia (32 from Southern China, 10 from Northern China, and 10 from Korea), 27 from Europe, 3 from the Near East, and 7 from Central Asia, as well as 6 *Sus cebifrons* individuals from Southeast Asia (Fig. 1a, Supplementary Table 2). This study generated a total of 4.82 × 10^10^ reads, with an average sequencing depth of 22.82× and a genome coverage of 98.73% (Supplementary Table 2). After aligning reads to the *Sus scrofa* 11.1 reference genome and applying stringent quality control filters, we identified 196,171,763 single-nucleotide polymorphisms (SNPs). Of these SNPs, 78.1 million are in intergenic regions, 86.9 million in intronic regions, and 5.7 million in exonic regions (Supplementary Table 3).

**Fig. 1.**
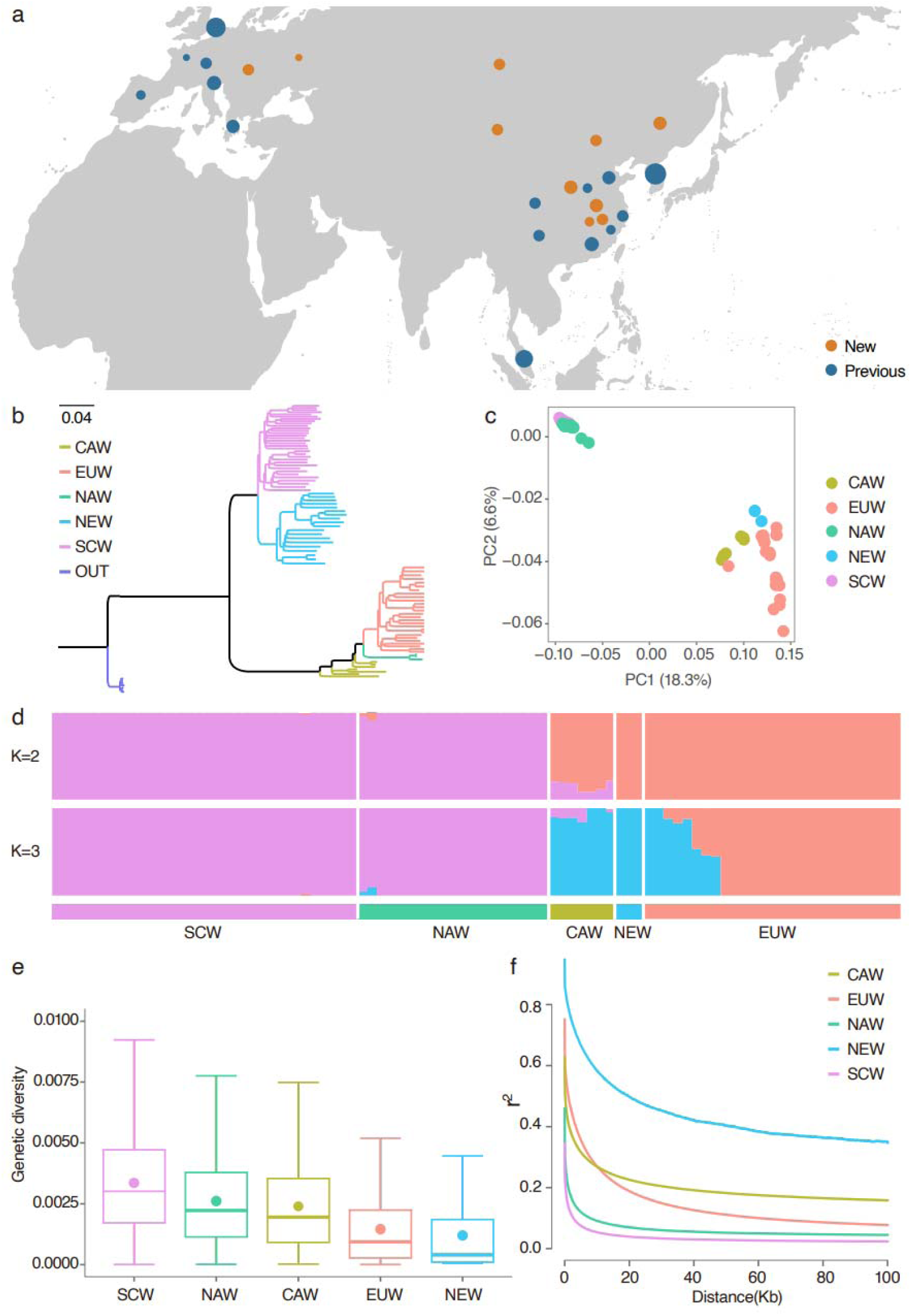
Geographic distribution and genetic structure of 89 wild boars. **a,** Worldwide distribution of studied wild boars. The point size indicates the population size. **b,** Neighbor-joining tree constructed using autosomal SNPs. SCW: Southern Chinese wild boars; NAW: Northeast Asian wild boars; CAW: Central Asian wild boars; EUW: European wild boars; NEW: Near Eastern wild boars; OUT: Outgroup (*Sus cebifrons*) **c,** Plots of principal components 1 and 2 for the 89 wild boars. **d,** Admixture pattern for K = 2 and K=3 reveals additional population structure among all 89 wild boars. The geographic locations are at the bottom. **e,** Genetic diversity of the different groups. **f,** Linkage disequilibrium patterns for the different groups.

### Genetic diversity and population structure

Both the phylogenetic tree (Fig. 1b) and the principal component analysis (PCA) (Fig. 1c; Supplementary Fig. S2) separated the wild boars into two clades, one including all East Asian wild boars (EAW) and the other including the Central Asian wild boars (CAW), European wild boars (EUW) and Near Eastern wild boars (NEW). Individual ancestry coefficients were inferred with Admixture to further assess population structure. With K = 2, which is also the optimal K-value (Supplementary Fig. S3), the ADMIXTURE analysis recapitulated a similar pattern as the PCA and phylogenetic tree. With K = 3, the third genetic ancestry (in yellow) appears mainly in Central Asian, Near Eastern and Greece wild boars (Fig. 1d).

Based on their geographical distribution, we subdivided the East Asian wild boars into the Southern Chinese wild boars (SCW) and the Northeast Asian wild boars (NAW, including individuals from Northeast China and Korea). Nucleotide diversity (π) analysis showed that the SCW had the highest levels of genetic diversity, followed by NEA, CAW, EUW, and NEW (Fig. 1e; Supplementary Fig. S4). This trend indicates a progressive reduction in genetic diversity moving from Southeast Asia towards Europe. The level of inbreeding measured by runs of homozygosity (ROH) was lower in EAW than in other groups (Supplementary Fig.S5). Genetic distances estimated via the pairwise fixation index (*F_ST_*) ranged from 0.112 (CAW-EUW) to 0.520 (NAW-NEW) (Supplementary Fig. S6). Additionally, linkage disequilibrium (LD) decay analysis revealed more rapid LD decay in the Southern Chinese population compared to other groups, suggesting its larger effective population size and higher genetic diversity over time (Fig. 1f). Overall, the observed genetic patterns reflect a clear geographic gradient of genetic variation, from Southeast Asia to Northern Asia, Central Asia, and finally Europe and the Near East, providing valuable insights into the spread of wild boars across Eurasia.

### Migration tracks and demographic history of the *Sus Scrofa*

To investigate the migration and expansion of *Sus scrofa* from Southeast Asia to Europe, we examined phylogenetic relationships and gene flow using TreeMix by using *Sus cebifrons* as the outgroup. The relationships inferred from the TreeMix analysis (Fig. 2a; Supplementary Fig. S7-8) are consistent with patterns observed in the phylogenetic relationships (Fig. 1c). Following the divergence between *Sus cebifrons* and *Sus scrofa*, the earliest split within *Sus scrofa* separated the CAW from the SCW. Subsequent branching involved the divergence of NAW from SCW, and EUW from CAW, with branch length reflecting the geographic distances from their origins in Southeast Asia. To infer demographic history of *Sus scrofa*, we applied Multiple Sequentially Markovian Coalescent (MSMC) analysis to 8 wild boar individuals, each with sequencing coverage exceeding 30×. The MSMC trajectories were largely consistent across individuals, indicating strong population cohesion throughout much of the species’ evolutionary history (Fig. 2b). The demographic history of *Sus scrofa* is marked by population fluctuations that correspond to glacial cycles during the Pleistocene. Around 600–800 thousand years ago (kya), all *Sus scrofa* populations experienced growth during the Pastonian interglacial stage, followed by a decline during the Mindel glaciation (455–300 kya), which persisted until the end of the Würm glaciation (110–10 kya).

**Fig. 2.**
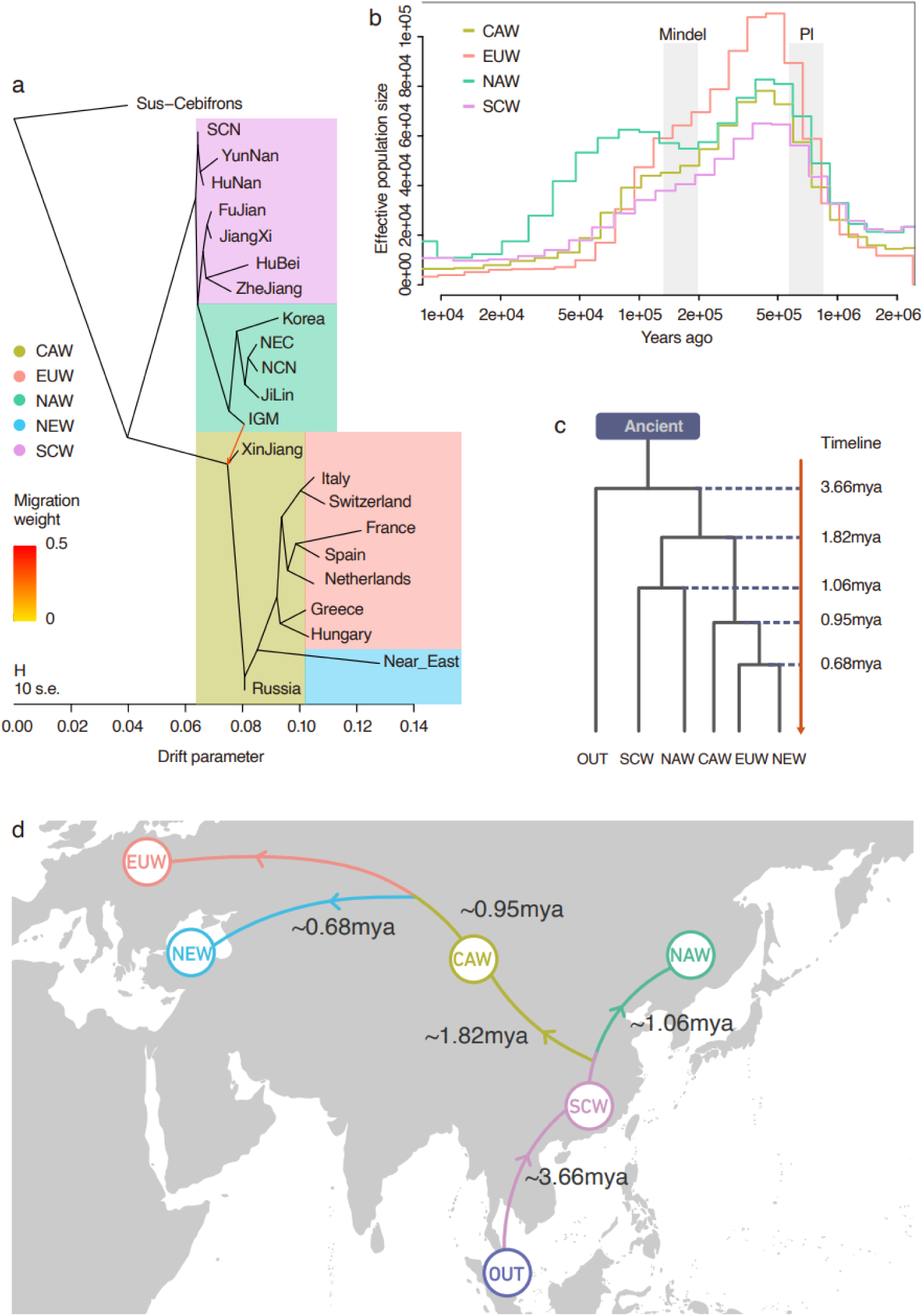
Demographic and migration histories for the wild boars. **a,** Tree topology inferred from TreeMix when one migratory tract is allowed. CAW: Central Asian wild boars; SCW: Southern Chinese wild boar; NAW: Northeast Asian wild boars; EUW: European wild boars; NEW; Near East wild boars. **b,** Demographic history of four geographical populations using MSMC. The gray shading marks the Pastonian interglacial stage (PI; 600-800 kya), the Mindel glaciation (455-300 kya), Würm glaciation (WG;110-10 kya). **c,** Inferred population demographic history of wild boars using the joint site frequency spectra. **d,** A proposed migratory history for wild boars from southern east Asia to Europe based on the evidence from our study.

To investigate population divergence, we performed a site frequency spectrum (SFS)-based analysis. First, we used SVDquartets to generate a phylogenetic tree, which was consistent with the relationships inferred from our TreeMix analysis (Supplementary Fig. S9). We then applied fastsimcoal2 to model demographic history such as divergence times, and migration rates between two groups that are geographically adjacent (Supplementary Fig. S10a), using SVDquartets inferred topology as input. Our analysis estimated that the divergence between *Sus cebifrons* and *Sus scrofa* occurred approximately 3.66 Mya. Following this, the ancestors of CAW/EUW/NE split off from the ancestor of SCW/NAW 1.8 Mya. At 1 mya the ancestors of the NAW branched off from SCW, migrating northeast into China and Korea. The lineages leading to the NEW and EUW separated from CAW approximately 0.95 million years ago, with the divergence between NEW and EUW occurring around 0.68 million years ago (Figs. 2c-d, Supplemental Fig. S10b).

The increased population size of EUW approximately 0.5 million years ago, as depicted in Fig. 2b, may suggest a population expansion in EUW following its divergence from NEW. A significant migration rate of 10.1% was identified between SCW and NAW, likely attributable to the minimal geographical barriers separating these two groups (Supplemental Fig. S10c). Notably, no significant population size differences were detected among *Sus scrofa* individuals (Supplemental Fig. S10d). Collectively, these findings elucidate a detailed timeline of population divergence events among *Sus* species and within *Sus scrofa* across Eurasia.

### Genomic signatures of selection on autosomes related to climatic adaptation

Originating in the wet and warm environments of Southeast Asia, *Sus scrofa* migrated across Eurasia and expanded into high-latitude regions generally characterized by colder climates. This migration likely imposed selective pressures that drove adaptation to diverse environmental conditions. To explore the genetic basis of these climatic adaptations, we conducted pairwise comparisons between CAW, which inhabit cold conditions, and SCW, which thrive in wet and warm environments (Supplementary Table 4). We performed window based (10kb) *F_ST_*, θπ ratio (θπ_2_/θπ_1_), and cross-population composite likelihood ratio (XP-CLR) analyses. By merging the top 5% of high-scoring regions from each analysis we identified 601 putative selected regions overlapping 320 genes (Fig. 3a, Supplementary Table 5-6). Kyoto Encyclopedia of Genes and Genomes (KEGG) pathway analysis revealed a significant over-representation of genes associated with metabolic pathways (Supplemental Fig. S11; Supplementary Table 7). These findings suggest that historical exposure to cold environments likely imposed strong selective pressures on basal metabolic rates, an important adaptation mechanism, given that most mammals increase basal metabolic activity in response to cold^23, 24^.

**Fig. 3.**
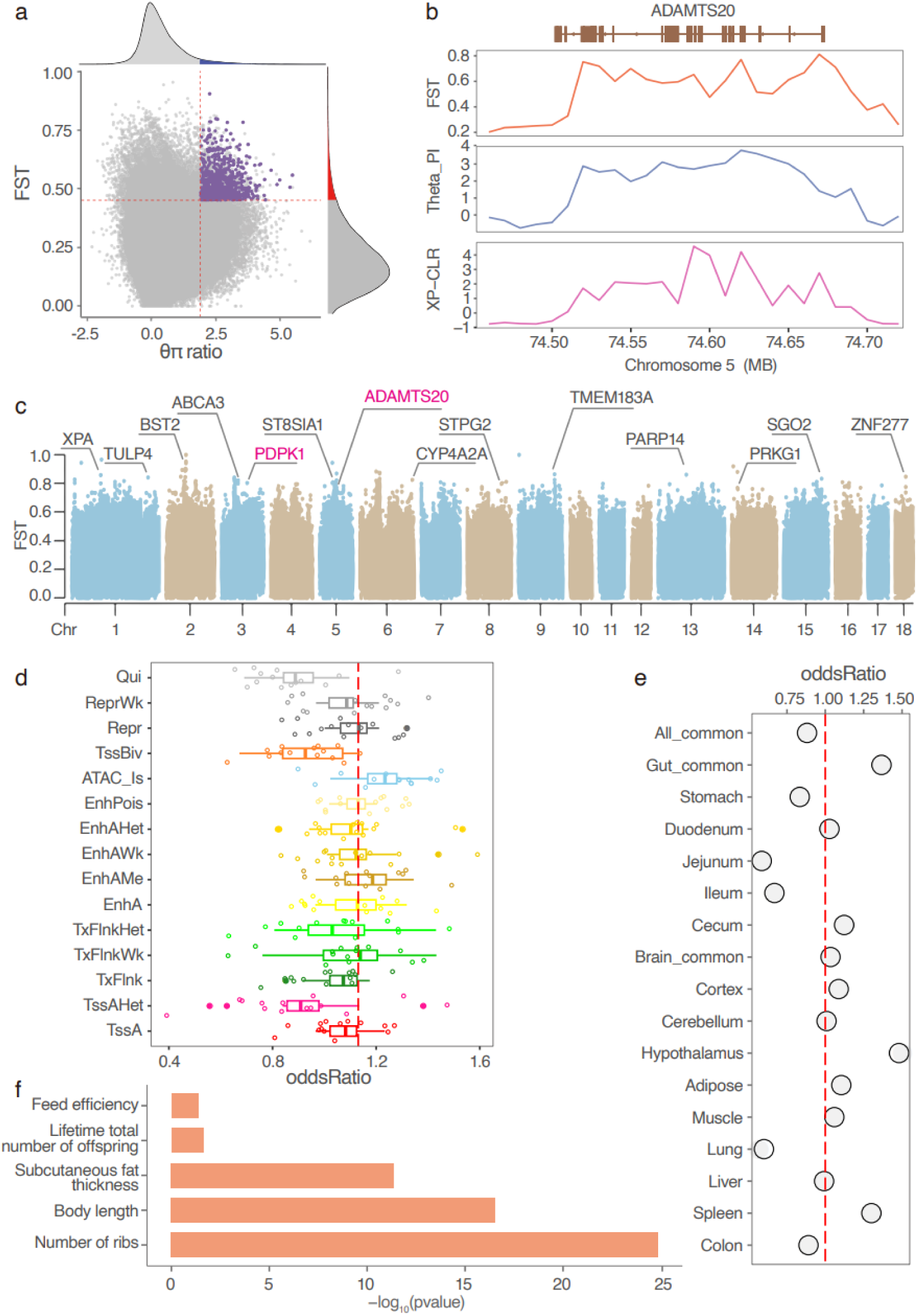
Genome-wide annotations of selective genome signatures for environmental adaptation. **a,** Distribution of the *F_ST_*, θπ ratio, and XP-CLR values for Central Asian wild boars (CAS) versus Southern China wild boars (SCN). The dashed vertical and horizontal lines indicate the significance thresholds for θπ ratio (1.7) and *F_ST_* (0.45) respectively. XP-CLR values largr than 1.9 were colored as blue. **b,** Strong selective signals on *ADAMTS20* based on *F*_ST_, θπ ratio, and XP-CLR for CAS versus SCN. **c,** Manhattan plots of the *F_ST_* (CAS vs. SCN). **d,** Selective genomic regions enrichment within 15 chromatin states. TssA: Strongly active promoters/transcripts; TssAHet: Flanking active TSS without ATAC; TxFlnk: Transcribed at gene; TxFlnkWk: Weak transcribed at gene; TxFlnkHe:Transcribed region without ATAC; EnhA:Strong active enhancer; EnhAMe:Medium enhancer with ATAC; EnhAWk:Weak active enhancer; EnhAHet:Active enhancer no ATAC; EnhPois:Poised enhancer; ATAC_Is:ATAC island; TssBiv:Bivalent/poised TSS; Repr:Repressed polycomb; ReprWk:Weak repressed polycomb; Qui:Quiescent. Values greater than 1 (dashed line) indicate significant enrichment. Each datapoint represents one of 14 different tissues. **e,** Selective genomic regions enrichment within tissue-specific enhancers. Values >1 (dashed line) indicate significant enrichment. All-common (shared among all tissues), gut-common (shared among gut tissues), and brain-common (shared among brain tissues). **f** Significantly enriched QTL terms for selective regions.

Among the identified genes, *ADAMTS20*, implicated in the regulation of melanocyte differentiation and pigmentation^25^, emerged as a key target of selection (Fig. 3b). Similarly, *PDPK1*, a central regulator of melanocyte proliferation—whose loss results in reduced pigmentation in mice^26^—was also identified as a gene under selection (Fig. 3c). Notably, *PDPK1* has been recognized for its role in local adaptation to skin pigmentation in African human populations^27^, further underscoring its evolutionary significance. The identification of *ADAMTS20* and *PDPK1* suggests that as wild boars expanded from lower-latitude to higher-latitude regions, these populations adapted to varying levels of ultraviolet (UV) radiation. In higher-latitude environments, where UV exposure is reduced, regulation of melanocyte differentiation and proliferation likely became crucial for maintaining adequate pigmentation, which in turn provides protection from environmental stresses. This highlights the adaptive importance of these genes in responding to environmental changes.

Our functional annotation analysis revealed that selective regions were predominantly enriched in open chromatin regions, with secondary enrichment in enhancer regions (Fig. 3d; Supplementary Table 8-9). This distribution suggests that adaptive evolution is more likely driven by regulatory changes in non-coding regions than by alterations in coding sequences. The significant concentration of selective sweeps in these regulatory regions underscores the critical role of gene regulation in enabling adaptive responses to environmental pressures.

Furthermore, our tissue-specific enhancer enrichment analysis identified brain, spleen, and gut as key organs for these regulatory adaptations (Fig. 3e). These findings suggest that these organs play critical roles in mediating adaptive responses: the brain, central to behavioral responses and environmental perception, adapts to new stimuli and conditions; the spleen, vital for immune function, often responds to new pathogens and immunological challenges; and the gut, essential for nutrient absorption and host-microbiome interactions, may adapt to dietary or ecological changes.

Our QTL (Quantitative Trait Loci) enrichment analyses reveal selective sweeps that are strongly linked to traits crucial for cold adaptation. These traits include variations in rib number, body length, and subcutaneous fat thickness—features that are integral to survival in colder climates (Fig. 3f; Supplementary Table 10). The observed increase in rib count and body length in certain populations may indicate structural adaptations that contribute to a more robust body form. Generally, mammals in warmer regions tend to be smaller than those in colder regions, likely due to the evolutionary advantage of a higher surface area-to-volume ratio, which facilitates more efficient heat dissipation^28^. Additionally, the enhanced subcutaneous fat thickness serves as a vital insulative layer, offering protection against extreme temperatures in high-latitude environments.

## Naturally selected variants for environmental adaptation

To detect selective single nucleotide polymorphisms (SNPs) associated with high-latitude adaptation, we employed the Cross-Population Extended Haplotype Homozygosity (xp-EHH) test, which identifies SNPs where a selected allele has reached, or is approaching, fixation in one population while remaining polymorphic in another. This analysis identified 18,582 variants as the top 1% loci, with 534 of these variants also detected by three other methods (*F_ST_*, θπ ratio, and XP-CLR). Consistent with the distribution shown in Fig. 3d, 49% of these shared variants (262 out of 534) are in regions of open chromatin, 24% (129 out of 534) are in enhancer regions, and only 5% are within gene bodies (Supplemental Table 11).

Among the 13 variants located in strong active enhancers, two were found to overlap with a muscle-specific enhancer in an intron of *LPIN1* (Fig. 4a; Supplementary Figure S12). Lipin-1 is a magnesium-dependent phosphatidic acid phosphohydrolase enzyme that plays a crucial role in triglyceride synthesis by catalyzing the dephosphorylation of phosphatidic acid to form diacylglycerol^29^. This enzyme is essential for adipocyte differentiation and also functions as a nuclear transcriptional coactivator, interacting with peroxisome proliferator-activated receptors to regulate the expression of genes involved in lipid metabolism^30^. In Lipin-1-deficient mice, the absence of this enzyme leads to lipodystrophy, characterized by underdeveloped adipocytes, impaired triglyceride utilization during fasting, and transient fatty liver and skeletal muscle myopathy due to disrupted glycerolipid homeostasis^31^. Haplotype analysis revealed that two variants, rs340542212 and rs324682561, are naturally selected in high-latitude populations, including NAW, CAW, and EUW (Fig. 4b). The ancestral alleles, rs340542212-C and rs324682561-C, have higher frequencies in low-latitude regions (0.85) compared to other populations (Fig. 4c; Supplemental Fig S13). *LPIN1* is highly expressed in adipose, testis, and muscle tissues (Fig. 4d), and the C allele is associated with lower expression of *LPIN1* in pig tissues, as indicated by PigGTEx data^32^ (Fig. 4e). Luciferase reporter assays conducted in pig primary myoblast cells showed that the ancestral C allele exhibits decreased enhancer activity compared to the derived T allele (Fig. 4f). These findings suggest that the rs340542212 and rs324682561 mutations in the muscle-specific enhancer influence triglyceride synthesis by modulating the enhancer activity and gene expression of *LPIN1*.

**Fig. 4.**
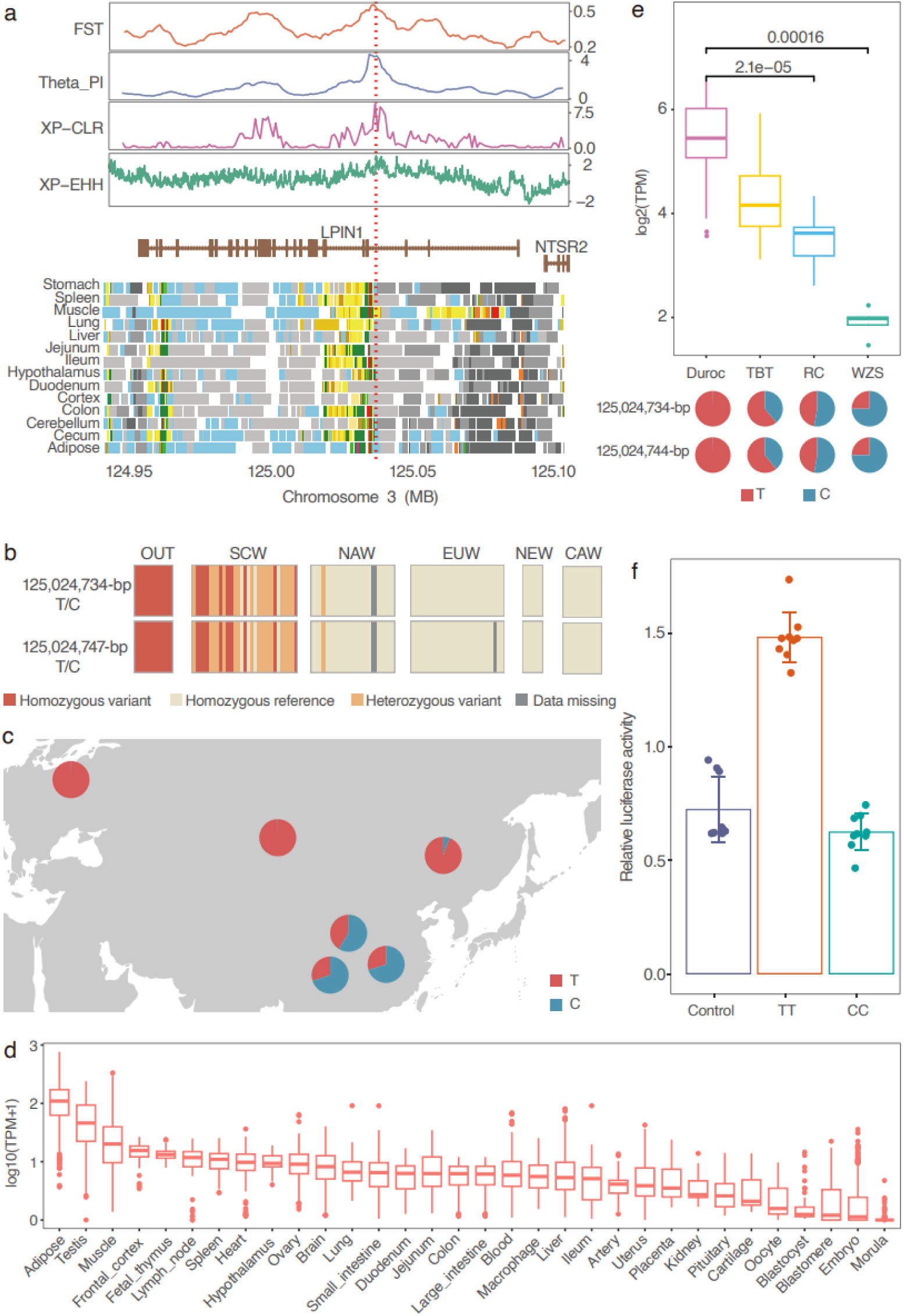
Two mutations affect the expression of *LPIN1* in skeletal muscle through regulating the enhancer activity. **a,** rs340542212 and rs324682561 overlap a muscle-specific enhancer (yellow) in the intron of *LPIN1*. Each color represents a specific chromatin state, with the corresponding relationships shown in Supplementary Figure S12. **b,** The patterns of genotypes of rs340542212 and rs324682561among the five wild boar groups and *Sus cebifrons* which were regarded as out group (OUT). CAW: Central Asian wild boars; SCW: Southern Chinese wild boar; NAW: Northeast Asian wild boars; EUW: European wild boars; NEW; Near East wild boars. **c,** Allele frequencies at rs340542212 in Eurasian populations located at different regions. **d,** Expression levels of *LPIN1* in different tissues. Expression levels are shown in log10(TPM). **e,** Expression levels of *LPIN1* in skeletal muscle tissue for different genotypes of rs340542212 and rs324682561 in domestic pigs. **f,** Luciferase reporter assay of rs340542212 and rs324682561 in skeletal muscle cells. N = 10.

We also identified 26 missense mutations among the 18,582 variants, spanning 9 genes (Supplementary Tables 12-13, Supplemental Fig S14-21). Notably, five missense mutations were found within the *ALPK2* gene, including one mutation, rs327271406, which has reached fixation in both CAW and EUW populations (Fig. 5a-b). Further analysis using data from the PigGTEX project confirmed that these mutations are also fixed in European domestic pigs (Fig. 5c). *ALPK2* is highly expressed in heart and muscle tissues (Fig. 5d), and phenome-wide association studies (phWAS)^33^ indicate a strong association between *ALPK2* and meat production traits (Fig. 5e; Supplemental Fig. S22; Supplemental Table 14). This finding is consistent with the established observation that European domestic pig breeds exhibit superior growth and meat production traits compared to their Asian counterparts, suggesting that the fixation of these mutations in *ALPK2* may contribute to these advantageous phenotypic traits.

**Fig. 5.**
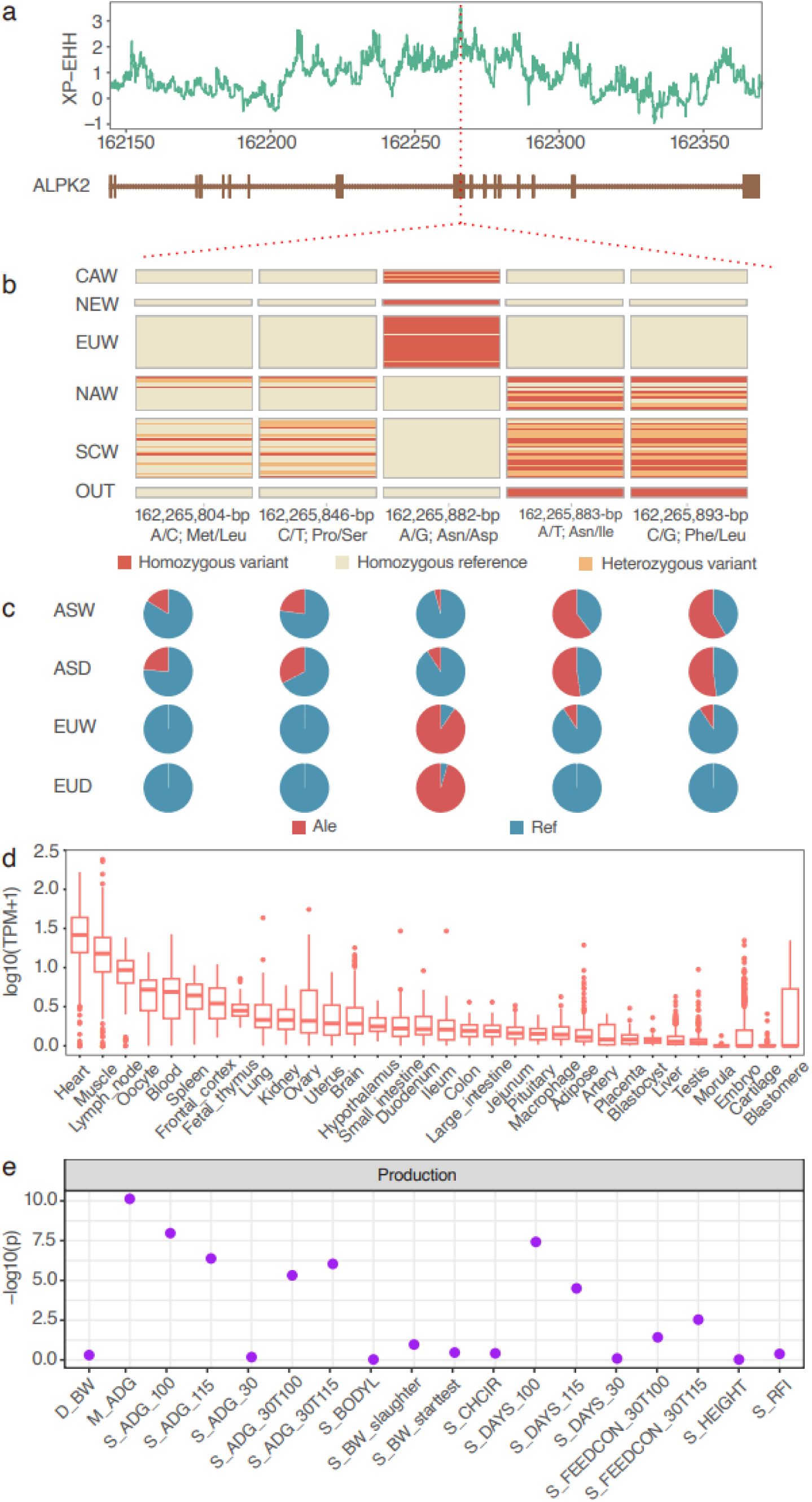
Genetic basis of ALPK2 for the meat production trait. **a** XP-EHH values around the *ALPK2* gene locus. **b** Haplotype pattern of the five 5 missense mutations within the *ALPK2* gene among the five wild boar groups and *Sus cebifrons* which were regarded as out group (OUT). CAW: Central Asian wild boars; SCW: Southern Chinese wild boar; NAW: Northeast Asian wild boars; EUW: European wild boars; NEW; Near East wild boars. **c** Allele frequency of the 5 missense mutation sites among pig populations in the pigGTEX project. ASW: Asian wild boars; ASD: Asian domestic pigs; EUW: European wild boars; EUD: European domestic pigs. **d** Expression levels of *ALPK2* in different tissues. Expression levels are shown in log10(TPM). **e** Point plot shows the association between ALPK2 and complex analysis. Data cite from PigBiobank^33^, each point represents the association between *ALPK2* and each trait in gene-based association analysis. The correspondence between the abbreviation and the full name of each trait is in the supplementary table 14.

## Discussion

In this study, we conducted an in-depth analysis of high-coverage whole-genome sequencing data from 89 wild boars, covering a broad geographic range from Southeast Asia to Europe, including previously underexplored regions in Central Asia. This comprehensive dataset allowed us to investigate the expansion routes of wild boars across Eurasia, shedding light on aspects of their dispersal that have remained elusive in previous research.

Earlier studies have made significant strides in tracing the migration patterns of wild boars and domestic pig, though certain details remain unresolved. For instance, Larson et al. traced the initial dispersal from Island Southeast Asia (ISEA) to the Indian subcontinent, followed by a gradual expansion into Eastern Asia and eventually reaching Western Europe^2, 12^. However, their work did not provide a detailed analysis of the specific migration pathways. Wu et al. later examined potential routes within East Asia^14^, but the relationships between these East Asian lineages and their western counterparts remained unclear. Additionally, a recent phylogeographic study using mitochondrial DNA and Y-chromosome data from wild boars proposed new hypotheses on the potential expansion routes across Asia, yet the historical connections between these lineages were not fully explored^22^.

Building on this foundation, our analysis provides a refined and more detailed picture of the evolutionary history of *Sus scrofa*. We estimate that the divergence between *Sus cebifrons* and *Sus scrofa* occurred approximately 3.6 million years ago which is consistent with previous studies^4, 5, 34^. Following the expansion from Southeast Asia into southern China, a significant bifurcation within *Sus scrofa* took place around 1.8 Mya, resulting in two distinct clades: SCW and CAW. About 1 Mya, the ancestors of NAW diverged from the Southern Chinese lineage and expanded northeastward into northeastern China and Korea. Subsequently, around 0.95 Mya, EUW and NEW diverged from the Central Asian populations and split at 0.68 Mya illustratingf the complex dynamics of population migration and adaptation within the *Sus scrofa* lineage.

By integrating high-coverage whole genome sequence data with historical genetic analyses, our study provides a more detailed and accurate reconstruction of ancient migration routes. While our findings partly align with previously proposed migration paths based on mitochondrial DNA and Y-chromosome data^22^. The lack of Indian samples restricts our ability to fully understand the complete range of dispersal routes and the potential interactions between different populations within Sus scrofa. Specifically, it may limit our understanding of how geographical barriers and ecological factors in South Asia influenced the evolutionary trajectories of wild boars. Furthermore, the genetic contribution of Indian wild boars to Eurasian populations remains unknown, leaving a gap in our comprehension of the species’ adaptive potential and population dynamics across diverse environments.

Previous studies on phenotype-associated variants in *Sus scrofa* have primarily focused on the effects of domestication and artificial selection on genomic variation^14, 35, 36, 37, 38^. In contrast, our study identified naturally selected loci that may be involved in phenotypic and physiological adaptation to diverse environments across wild boar populations. Similar to the domestication process, we found that climate adaptation was predominantly influenced by mutations in regulatory regions rather than coding regions^39, 40^. Gene functional enrichment analysis revealed that metabolic processes were the primary functional categories associated with cold adaptation, supporting the idea that energy regulation and metabolic efficiency are crucial for survival in cold climates.

Moreover, the functions of naturally selected genes showing significant differences in non-synonymous SNP allele frequencies between CAW and SCW populations provided further evidence for selection signatures on genes related to biological processes such as cellular signaling, metabolism, endocrine function, protein kinase activity, transporter functions, and development. These findings indicate that climate-driven adaptation in *Sus scrofa* has shaped a variety of physiological and biochemical pathways, highlighting the intricate relationship between genetic variation and environmental pressures.

In summary, our study provides a refined understanding of *Sus scrofa*’s migration and adaptation, revealing detailed expansion routes and climate-driven genetic changes. The findings emphasize the role of regulatory regions in adaptation and highlight metabolic pathways as key factors in cold tolerance. By integrating high-resolution genomic data, we advance the understanding of wild boar evolutionary history and set the stage for future research on local adaptation and environmental impacts on mammalian evolution.

## Methods

### Sample collection and sequencing

Samples from individuals with information about geographic origin were collected. Total genomic DNA was extracted from ear tissues using standard phenol/chloroform methods. For each individual, ∼3 μg DNA was sheared into fragments of 500 bp with the Covaris system. DNA fragments were then processed and sequenced using the Illumina HiSeq 2500 platforms. Furthermore, published genomic data for 72 wild boars were downloaded from NCBI. Raw reads were filtered with the following criteria: reads with unidentified nucleotides exceeded 10% were discarded and reads with the proportion of low-quality base (Phred score ≤ 5) larger than 50% were discarded.

### Read Mapping and SNP Calling

We processed and analyzed the whole genome sequence data using the following pipeline. Briefly, we trimmed the raw sequence reads by Trimmomatic (v0.39)^41^, with the default parameters. Both low-quality bases and adapter sequences were removed to ensure high-quality data for downstream analysis. Clean reads were mapped to Sscrofa11.1 using BWA-MEM (v0.7.5a-r405) with default parameters^42^. We marked duplicated reads by Picard (v2.21.2) (http://broadinstitute.github.io/picard/). We removed 16 samples with low read depth (< 5x). Finally, we kept 108 samples for jointly calling variants using the Genome Analysis Toolkit (GATK) (v4.1.4.1)^43^ with parameters: QD> 2, MQ < 40, FS > 60, SOR > 3, MQRankSum < −12.5 and ReadPosRankSum < −8, yielding ∼214 million SNPs. We removed SNPs with MAF < 0.01 and/or missing call rate > 0.9 using bcftools (v1.9)^44^ and employed Beagle (v5.1)^45^ to phase the filtered variants and impute sporadically missing genotypes. Finally, a total of 42,523,218 SNPs were retained. These SNPs were annotated by SnpEff v.4.3^46^.

### Sample filtering, Genetic diversity and Population Structure

We obtained 2,573,646 linkage disequilibrium (LD) independent SNPs using PLINK (v1.90) with parameters: --indep pairwise 1000 100 0.2^47^. We calculated the IBS matrix among the 108 samples using GenABEL^48^ package in R. Then, 98 unrelated wild boars were selected from 108 samples by removing samples that have a common ancestor within 3 generations. We used all high-quality SNPs to calculate nucleotide diversity (π) and population genetic differentiation (*F_ST_*) with 10-kb nonoverlapping windows using vcftools (v1.2.6)^49^.

The principal components analysis (PCA) was conducted using PLINK (v1.90b4.4)^47^. A neighbor-joining phylogenetic tree was constructed according to the P-distance using TreeBest (v1.9.2)^50^, with a bootstrap value of 1,000 to elucidate phylogenetic relationships from a genome-wide perspective. The population structure was analyzed using the expectation–maximization algorithm in ADMIXTURE (v1.3.0)^51^, with the ancestry-specifying K ranging from 2 to 10 and 10,000 iterations per run. LD was calculated using PopLDdecay (v3.30)^52^.

### Effective population size and divergence time inference

We estimated effective population sizes using Multiple Sequentially Markovian Coalescent (MSMC2)^53^ with -p ‘4+50*1+4+6’^54^. The porcine generation time (g) = 5 years, and neutral mutation rate per generation (μ) = 2.5 × 10^−8^ were based on previous reports^16^.

We used Treemix v1.13-7^55^ to test models of possible admixture 89 wild boars with *Sus cebifrons* used as outgroup. 2,573,646 LD independent SNPs were used as input. Migrations from m0 to m9 were tested, with 10 replicates per m to assess consistency.

We used PAUP∗version 5.0 to run SVDquartets^56^ to estimate the branching pattern among the five populations (SAW, SCW, NAW, CAW, and EUW). SNPs were filtered by bcftools with the following parameter: -e ‘AC==0 || AC==AN || F_MISSING > 0.2’ -m2 -M2^57^. A Ruby script (https://github.com/mmatschiner/tutorials/blob/master/species_tree_infere nce_with_snp_data/src/convert_vcf_to_nexus.rb) was used to convert the VCF file into PAUP* input file.

The joint SFS approach implemented in fastsimcoal2^58^ was performed to model more recent demographic fluctuations and respective divergence times based on the species tree estimation by SVDquartets. To mitigate the effect of LD, we filtered out the SNPs located within the 15 chrome states regions. The multidimensional folded SFS for all five populations was generated with easySFS. The conditional maximization algorithm (ECM) was used to maximize the likelihood of each parameter while keeping the others stabilized. This ECM procedure runs through 40 cycles where each composite likelihood was calculated using 100,000 coalescent simulations. The best parameters were determined according to the maximum value of the likelihoods and Akaike information criterion.

### Selective signatures Analysis

To identify genomic regions that may have been subject to selection for different environment adaptation, we scanned the genome comparisons between Central Asian and Southern Chinese wild boars. We calculated θπ for each population and the *F_ST_* between the two populations using VCFtools with a genomewide sliding window strategy (10 kb in length with 10-kb step)^49^. The log value of θπ ratio (θπ_SCN_/θπ_CAS_) was estimated. Multilocus allele frequency differentiation based selective sweeps (XP-CLR) were detected using xpclr tool^59^ with a 10kb sliding window. Candidate regions under positive selection were extracted based on the top 5% value of *F_ST_*, log (θπ ratio) and XPCLR.

We used cross-population extended haplotype heterozygosity test (XP-EHH)^60^ to detect the SNPs that have reached or are approaching fixation in one population, while remaining polymorphic in another. The top 1% outliers of all SNPs were selected as candidate selective variances.

### Functional enrichment analysis of selective signatures

We pinpointed genes located in the candidate selective regions based on the Sscrofa11.1 reference genome assembly. Genes that overlapped with these regions were selected for further analysis. Kyoto Encyclopedia of Genes and Genomes (KEGG) enrichment analysis of these genes was implemented by using the Database for Annotation, Visualization and Integrated Discovery (DAVID)^61^.

We downloaded from publicly available datasets^62^. These datasets describe the genomic features including promoters (TssA, TssAHet, and TssBiv), TSS-proximal transcribed regions (TxFlnk, TxFlnkWk, and TxFlnkHet), enhancers (EnhA, EnhAMe, EnhAWk, EnhAHet, and EnhPois), repressed regions (Repr and ReprWk), and quiescent regions (Qui) in a BED file for each tissue. The significance of enrichment analysis for candidate selective regions were performed by using the R package LOLA v1.32.0^63^ with 15 chromatin states for 14 different pig tissues used as reference annotation dataset. We used R package GALLO v1.4^64^ for QTL enrichment analyses against the pig QTL Database, and items with P-values under 0.05 from multiple tests were considered as enriched ones.

### Isolation of pig primary myoblast and cell culture

Pig primary myoblasts were isolated from a 2-day-old Duroc pig as described previously^65^ and cultured in DMEM/F12 -Dulbecco’s Modified Eagle Medium (DMEM/F12, Gibco, USA) supplemented with 10% fetal bovine serum and 1% penicillin-streptomycin (PS, Thermo Scientific, USA) at 37 in a 5% CO2 incubator.

### Functional analyses of rs340542212 and rs324682561 at the LPIN1 locus

A 51 bp sequence containing rs340542212 and rs324682561 with 25 bp up and 25 downstream of rs340542212 was synthesis and cloned into the pGL4.23-basic Luciferase Reporter Vector (GeneCreate, China). Candidate functional mutations of rs340542212 and rs324682561 were synthesized as well. Upon reaching approximately 60% confluency, the Pig primary myoblasts were transfected with plasmids. The luciferase vector containing the SNP site (experimental vector; 1 μg) and the control reporter vector pRL-TK (20 ng) were co-transfected into the Pig primary myoblasts at a 50:1 ratio using Attractene Transfection Reagent (Qiagen, Germany) according to the manufacturer’s protocol. After 24 h of transfection, the transfected cells were collected and lysed. Luciferase activity was then measured using the Dual-Luciferase Reporter Assay System (Promega, USA), and the results were normalized to Renilla luciferase activity.

## Data availability

All raw data analyzed in this study are publicly available for download without restrictions from SRA (https://www.ncbi.nlm.nih.gov/sra/) databases. Details of Whole Genome Sequenceing datasets can be found in Supplementary Tables 1. All the WGS data newly generated in this study are available under CNCB GSA (https://ngdc.cncb.ac.cn/) accessions PRJCA016120.

## Declaration of competing interest

The authors declare that they have no conflict of interest.

## Acknowledgements

This work was supported by grants from the Shenzhen Outstanding Talents Training Fund (Grand NO. 202102), the National Key R&D Program of China (Grant NO. 2021Y FD301201, 2024YFF0728800), the National Natural Science Foundation of China (Grant NO. 31972539, 31501933, 32102513), the STI 2030 — Major Projects (Grant NO. 2022ZD04017), the National Natural Science Foundation of Guangxi province (Grant NO.2024GXNSFAAO10105), the Science, Technology, and Innovation Commission of Shenzhen Municipality (Grant NO. JCYJ20180306173644635), Science Technology Innovation and Industrial Development of Shenzhen Dapeng New District (Grant NO.PT20170201, PT202101-05), the China Scholarship Council (Grant NO.202003250046).

## Notes

### Competing Interest Statement

The authors have declared no competing interest.

