## Supplementary figures for "Genomic Insights into the Demographic History and Local Adaptation of Wild Boars Across Eurasia"

This supplementary file contains supplementary Figures 1-22 and supplementary references

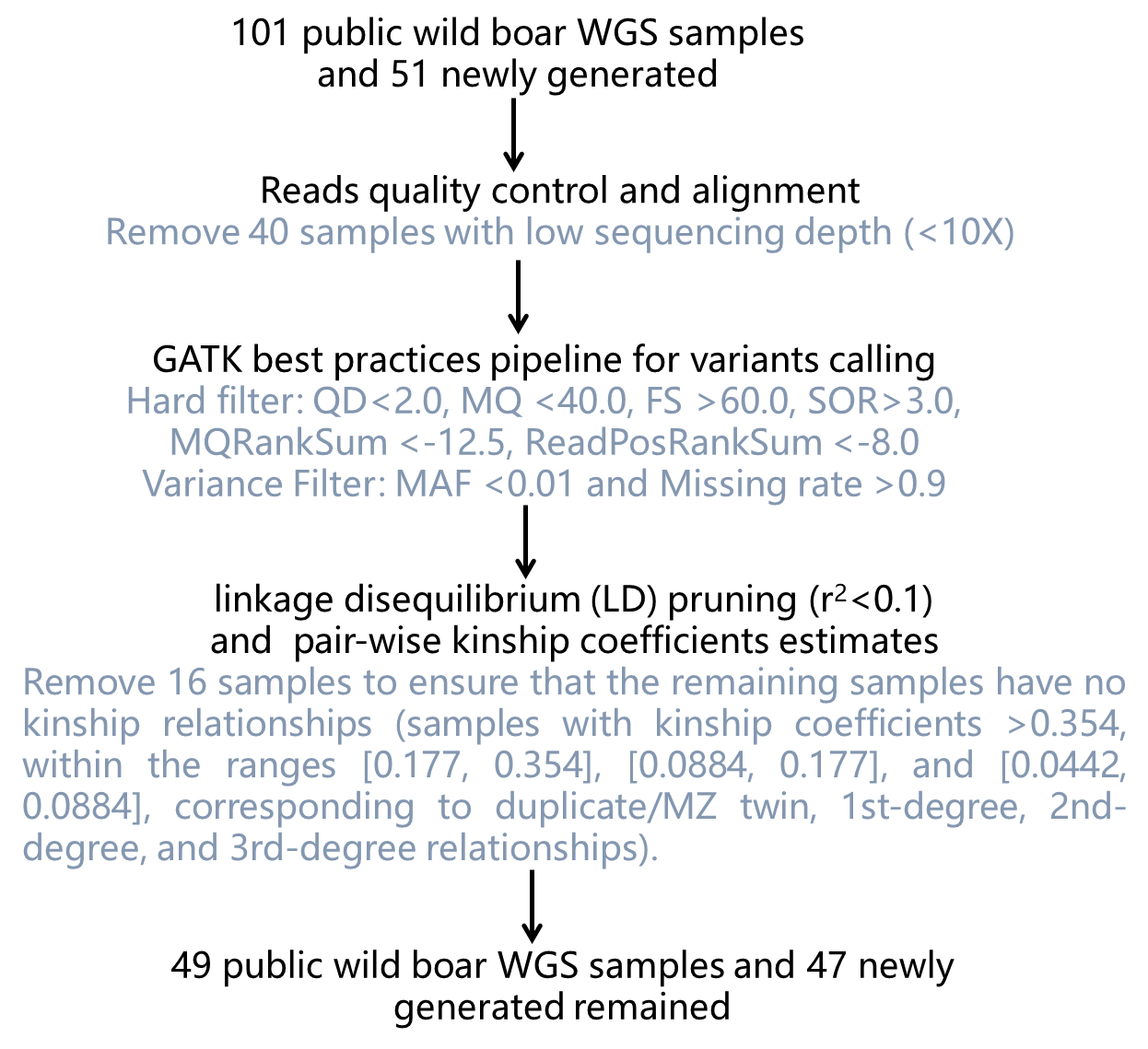

**Figure S1 The work pipeline utilized to filter samples and SNPs from 154 WGS individuals.**

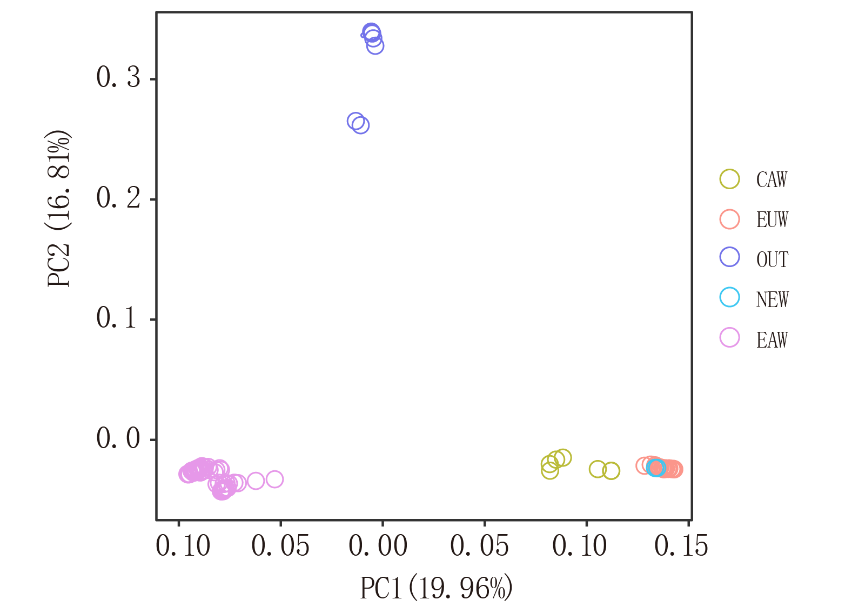

**Figure S2 Principal component analysis (PCA) of all 154 samples, illustrated by PC1 against PC2. Colors reflect the geographic regions of sampling. OUT: *Sus cebifrons* which were regarded as out group; EAW: East Asian wild boars; CAW: Central Asian wild boars; EUW: European wild boars; NEW: Near Eastern wild boars.**

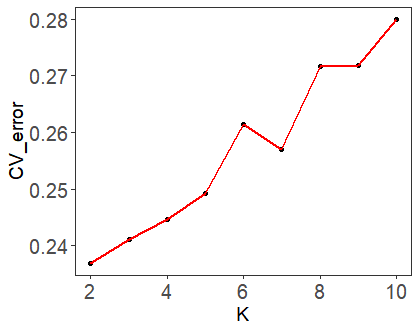

**Figure S3 Cross-validation (CV) error values of each K from 2 to 10**

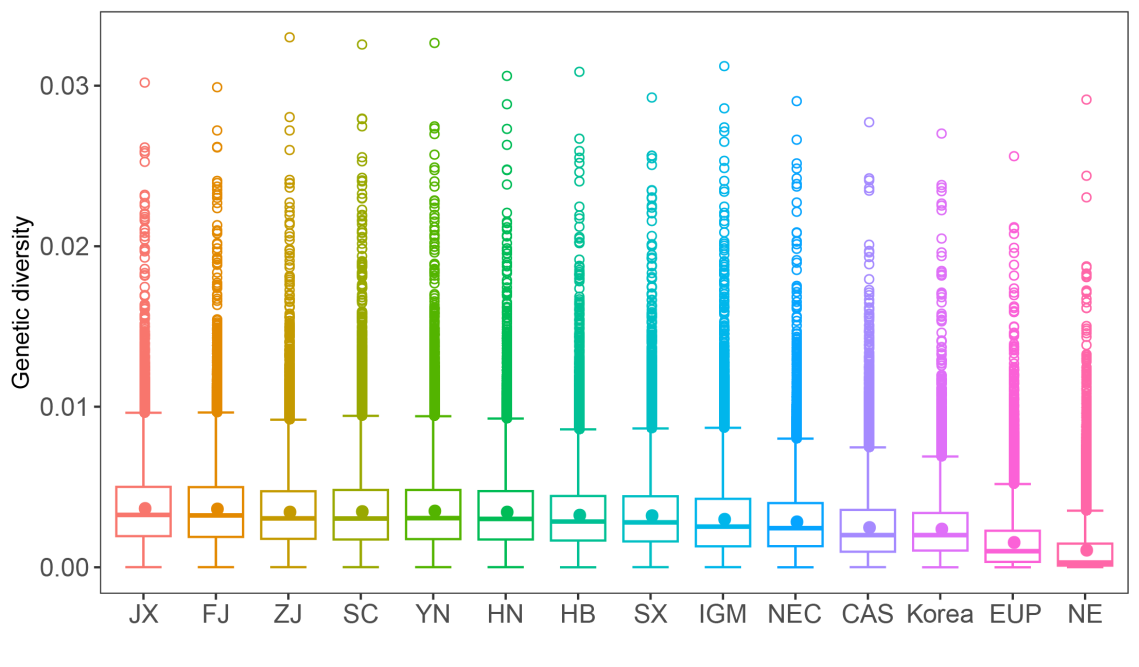

**Figure S4 Genetic diversity for the 89 wild boars. X-axis show the region where the wild boars come from. JX, Jiangxi province; FJ, Fujian province; ZJ, Zhejiang province; SC, Sichuan province; YN, Yunnan province; HN, Hunan province; HB, Hubei province; SX, Shanxi province; IGM, Inner Mongolia; NEC, Northeast China (including Heilongjiang and Jilin province); CAS, Central Asia; EUP, Europe; NE, Near East.**

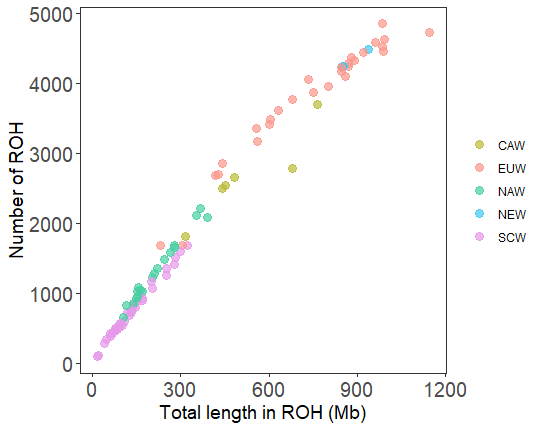

**Figure S5 Distribution pattern of runs of homozygosity (ROH) in 89 wild boars. Total autosomal SNPs were used for ROH analysis using PLINK. The final parameters were set to a minimum length of 100 kb, a scanning window size of 50 SNPs, a minimum density threshold of 200 SNPs, a large gap of 1000 kb, a maximum number of heterozygous SNPs in the scanning window of 1, and a scanning window threshold level of 0.05. The results show that these settings yield the expected number (maximum number was 4,848) and total length (maximum length is 1,145.81 Mb) of ROH. (SCW: Southern Chinese wild boars; NAW: Northeast Asian wild boars; CAW: Central Asian wild boars; EUW: European wild boars; NEW: Near Eastern wild boars).**

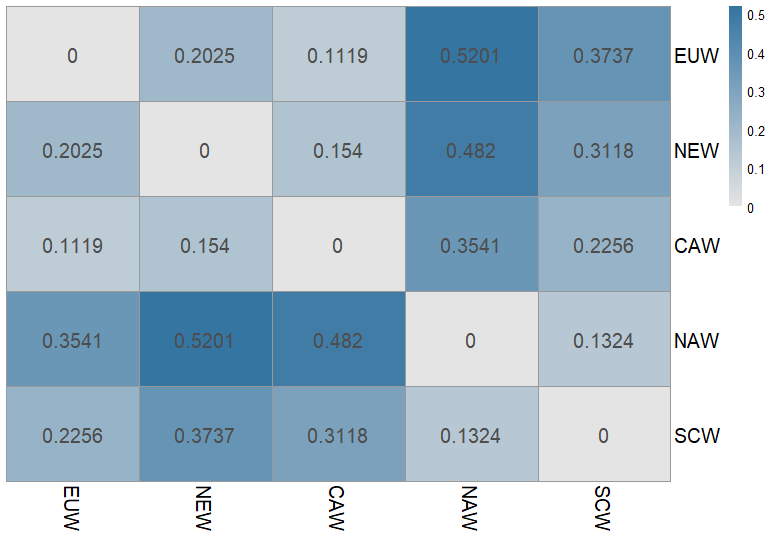

**Figure S6 Mean pairwise *F_ST_* values between each group of wild boars. A total of 89 samples and autosomal SNPs were used for *F_ST_* analysis using vcftools. (SCW: Southern Chinese wild boars; NAW: Northeast Asian wild boars; CAW: Central Asian wild boars; EUW: European wild boars; NEW: Near Eastern wild boars).**

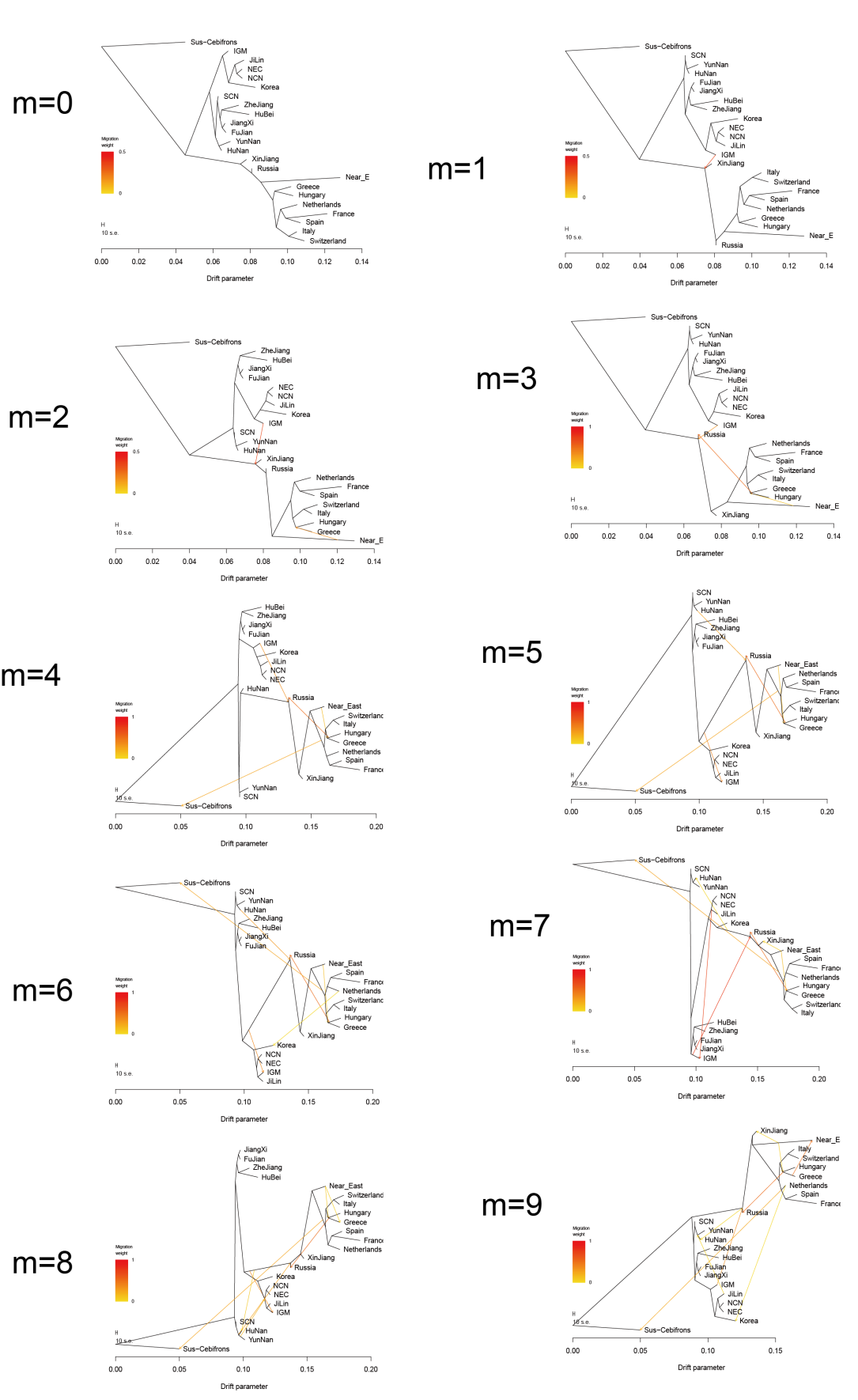

**Figure S7 Inferences of population splits and admixture using TreeMix. Tree and migration edges for 0 to 9 migration edges.**

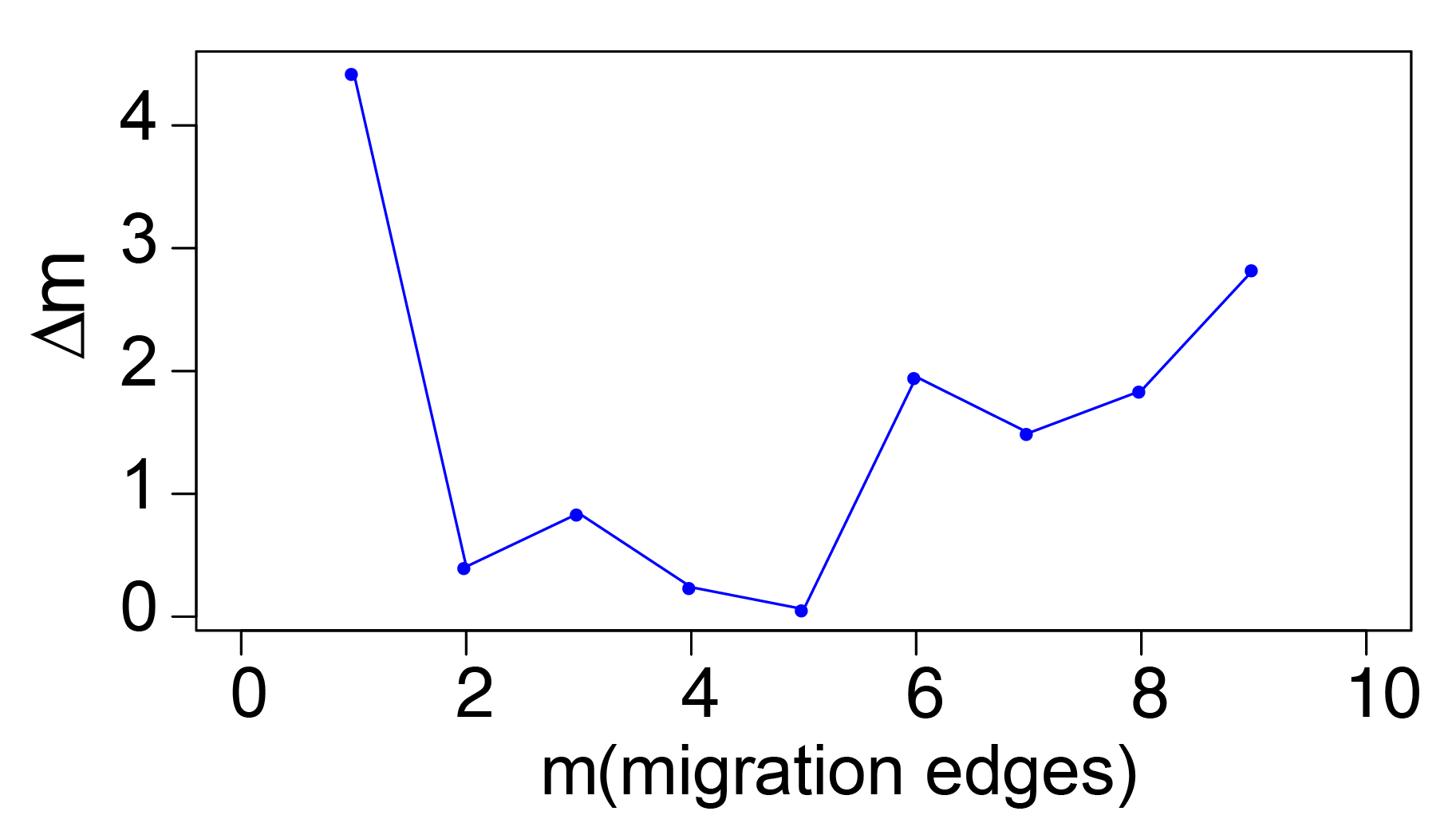

**Figure S8 Variation explanation change for adding an additional migration edge from 1 to 9. The output produced by OptM. The peak in second-order rate of change (Δ*m*) across values of m at 1 edge.**

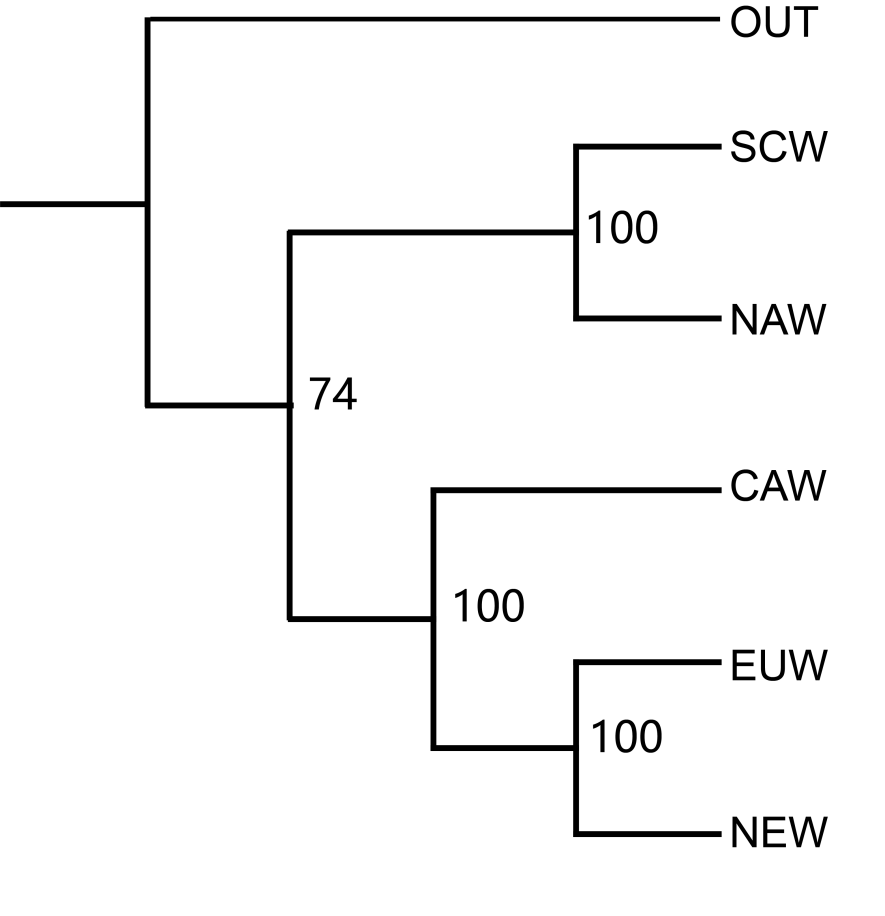

**Figure S9 The species tree based on SVDquartets+PAUP*. The numbers represent node support inferred from 100 non-parametric bootstrap repetitions.**

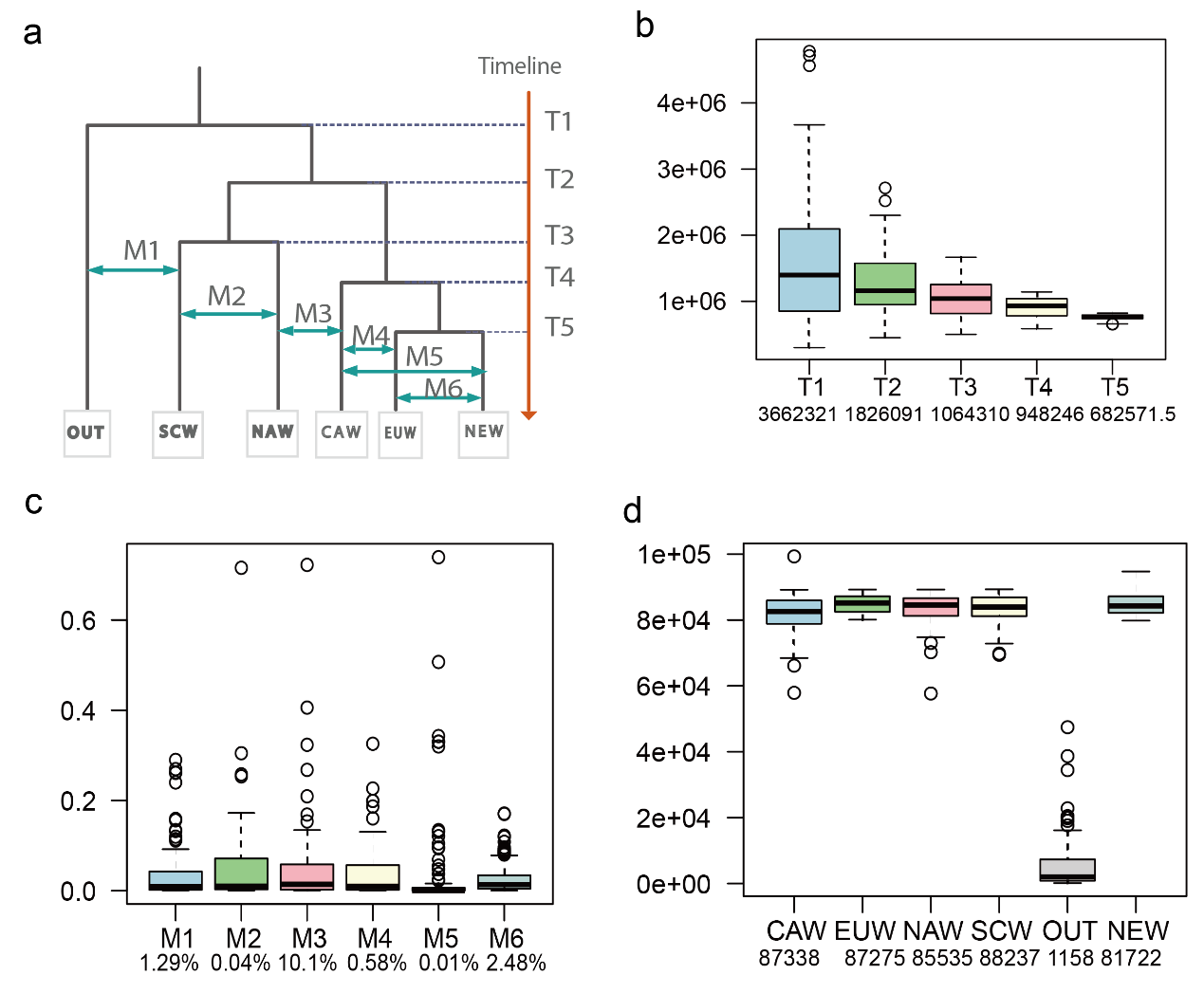

**Figure S10 Confidence Intervals from 100 parametric bootstraps for inferred**

**demographic parameters.** (a) The demographic model used for fastsimcoal2 analyses and parameters evaluated. T1-T6 represent the divergence time. Two-way arrows indicate the migration rate. Light gray squares at the bottom represent the effective population sizes of each group. (b) Divergence: the divergence time of each group (all times were converted to years assuming a generation time of 6.5 years). (c) Migration rate: proportion of individuals that move from one population to another per generation. (d) The effective population sizes of each group.

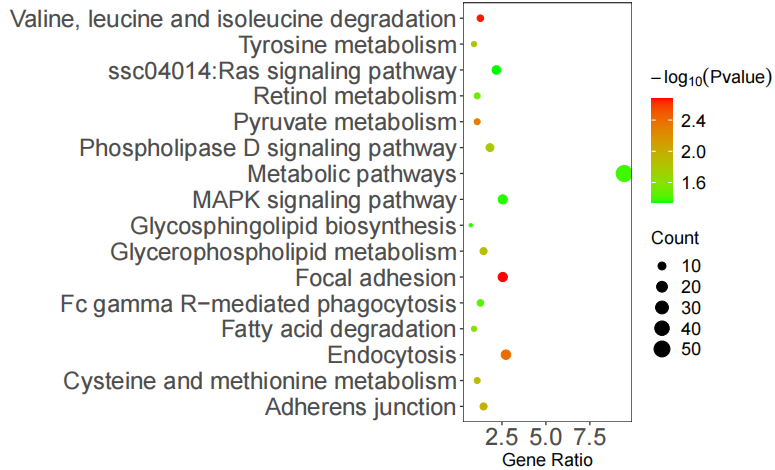

**Figure S11 Significant KEGG pathways for 320 genes identified in Fig.3a. The color of the circle represents the −log_10_ (*p* value). The size of the circle represents the number of selected genes in the enriched pathway.**

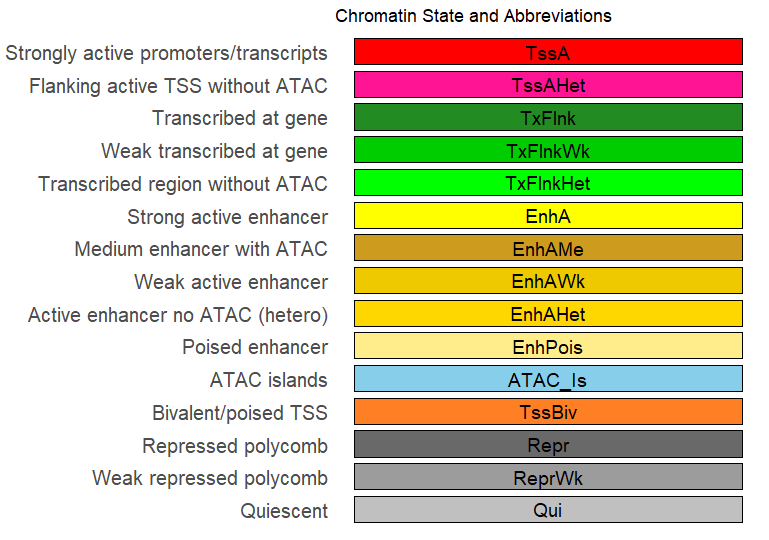

**Figure S12 Definitions and abbreviations of 15 chromatin states detected previously^1^ from 14 pig tissues, including stomach, spleen, muscle, lung, liver, jejunum, ileum,hypothalamus, duodenum, cortex, colon, cerebellum, cecum, and adipose. In the lower part of Fig. 4a, colors indicate the corresponding chromatin states.**

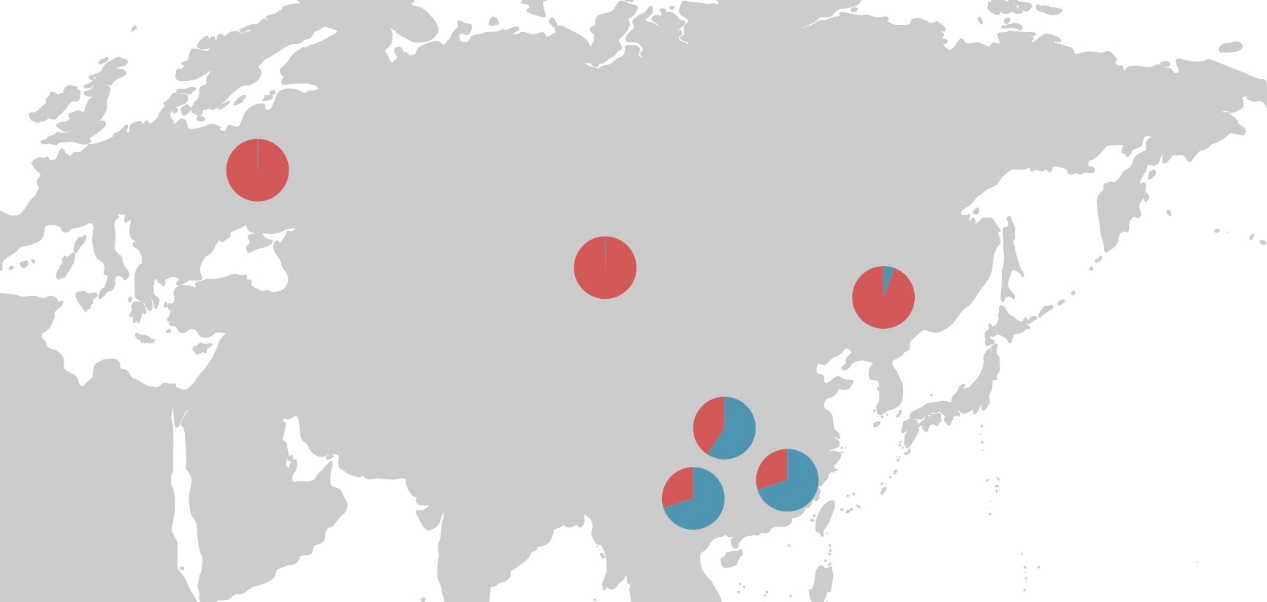

**Figure S13 Allele frequencies at rs324682561in Eurasian populations located at different regions**

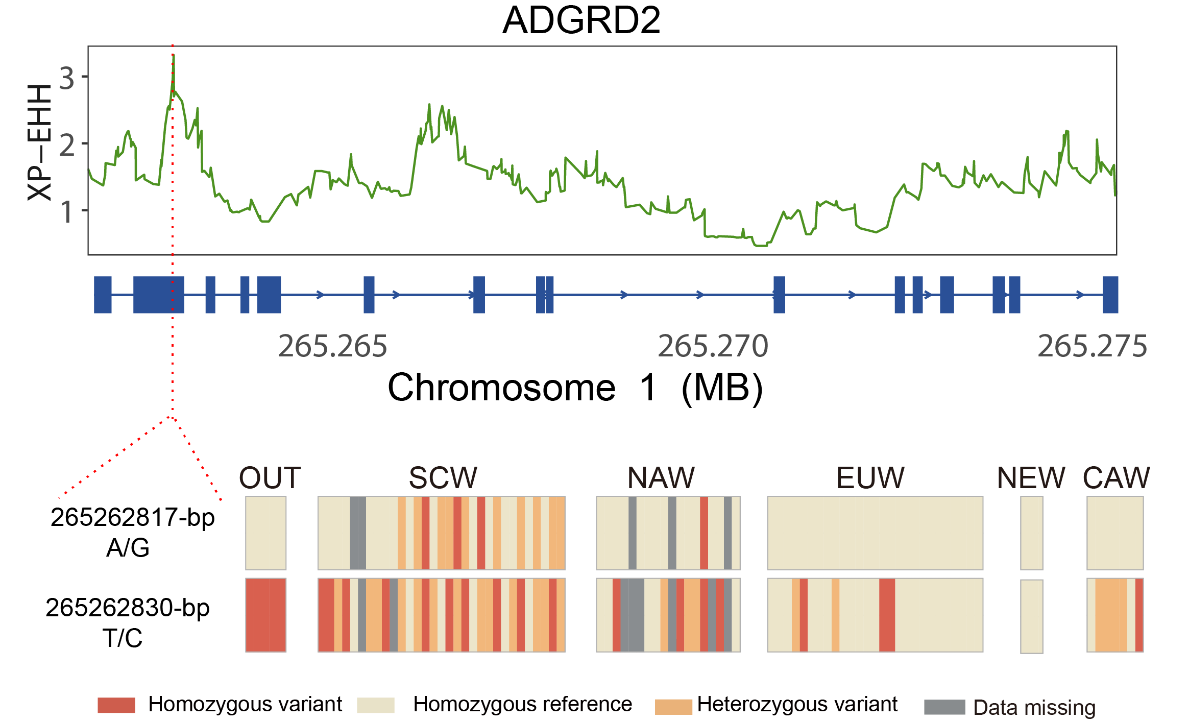

**Figure S14 XP-EHH values around the *ADGRD2* gene locus and haplotype pattern of the two missense mutations within the *ADGRD2* gene among the OUT and 89 wild boars. OUT: *Sus cebifrons* which were regarded as out group; SCW: Southern Chinese wild boars; NAW: Northeast Asian wild boars; CAW: Central Asian wild boars; EUW: European wild boars; NEW: Near Eastern wild boars.**

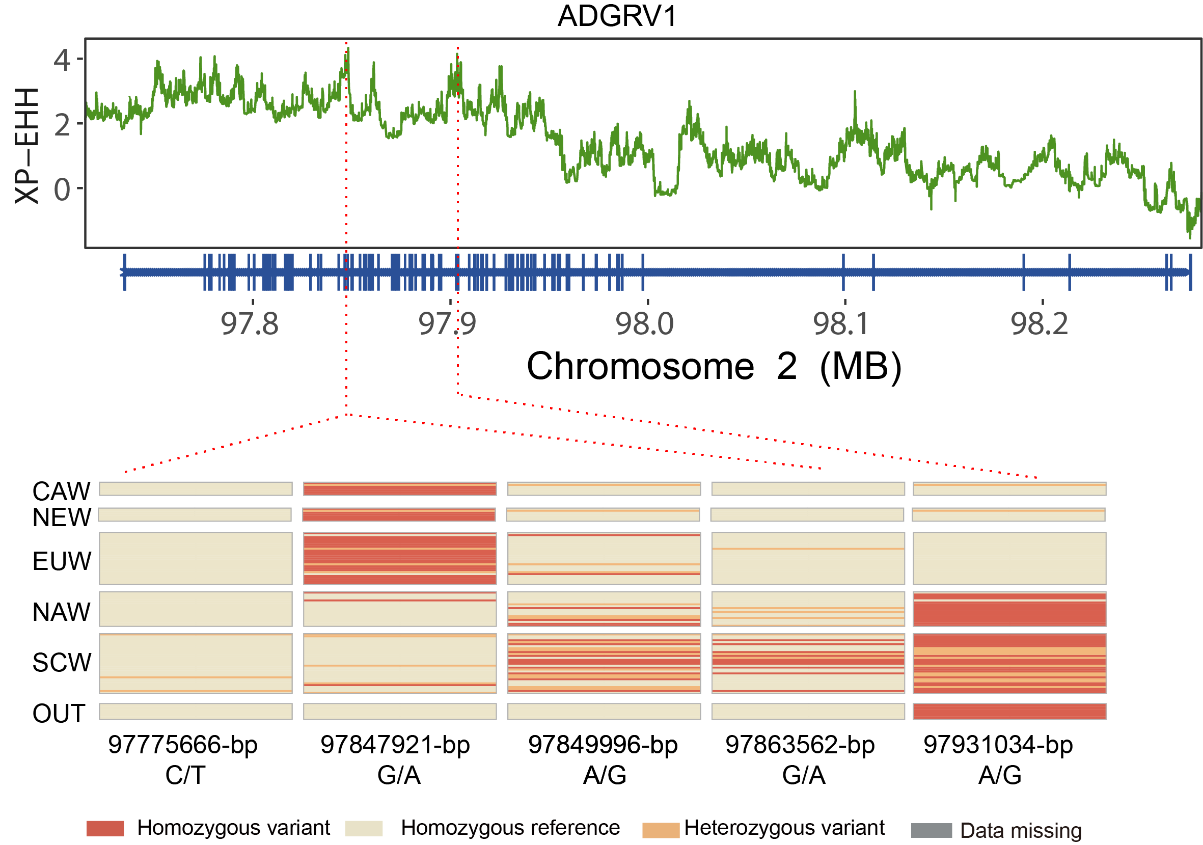

**Figure S15 XP-EHH values around the *ADGRV1* gene locus and haplotype pattern of the five missense mutations within the *ADGRV1* gene among the OUT and 89 wild boars. OUT: *Sus cebifrons* which were regarded as out group; SCW: Southern Chinese wild boars; NAW: Northeast Asian wild boars; CAW: Central Asian wild boars; EUW: European wild boars; NEW: Near Eastern wild boars.**

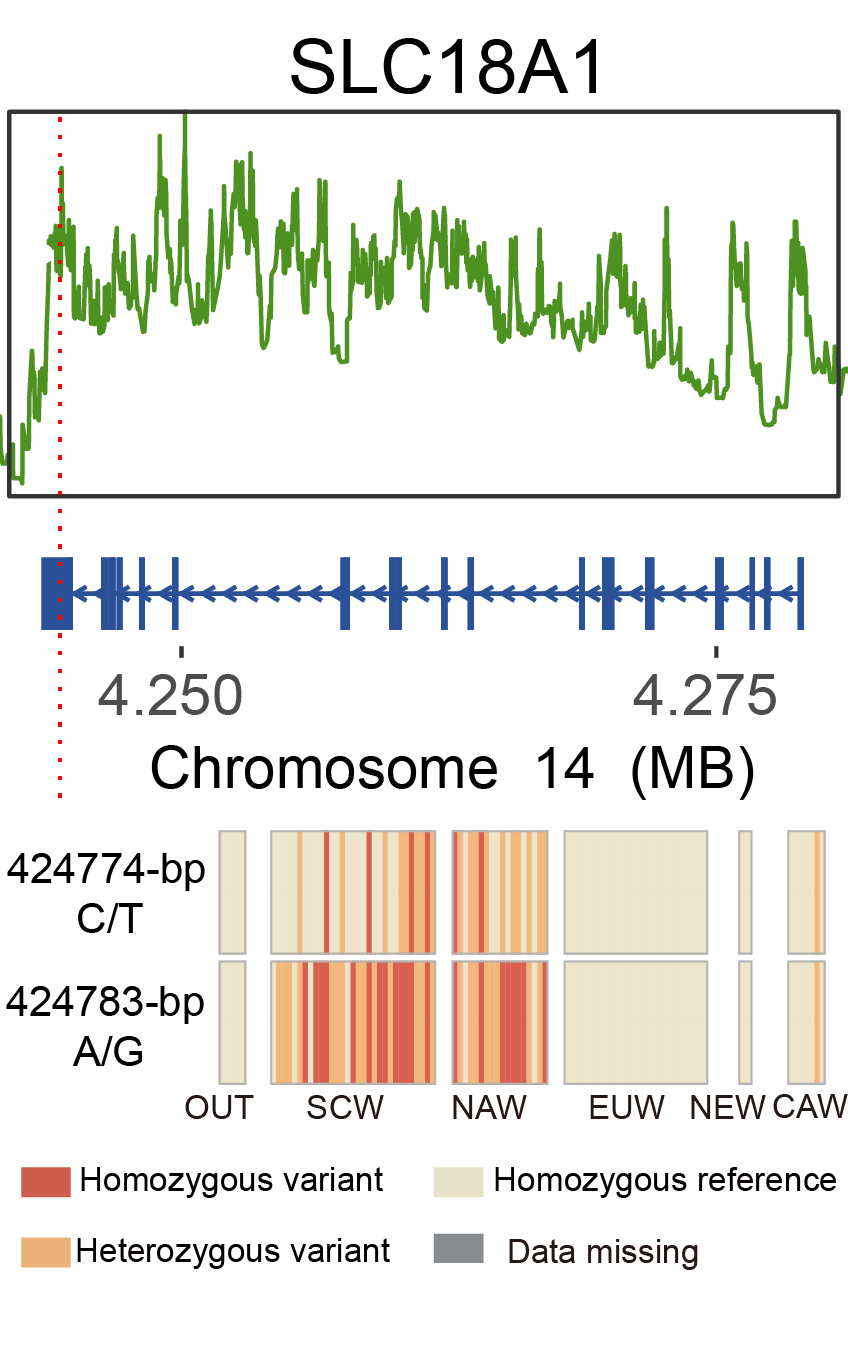

**Figure S16 XP-EHH values around the gene locus and haplotype pattern of the two missense mutations within the *SLC18A1* gene among the OUT and 89 wild boars. OUT: *Sus cebifrons* which were regarded as out group; SCW: Southern Chinese wild boars; NAW: Northeast Asian wild boars; CAW: Central Asian wild boars; EUW: European wild boars; NEW: Near Eastern wild boars.**

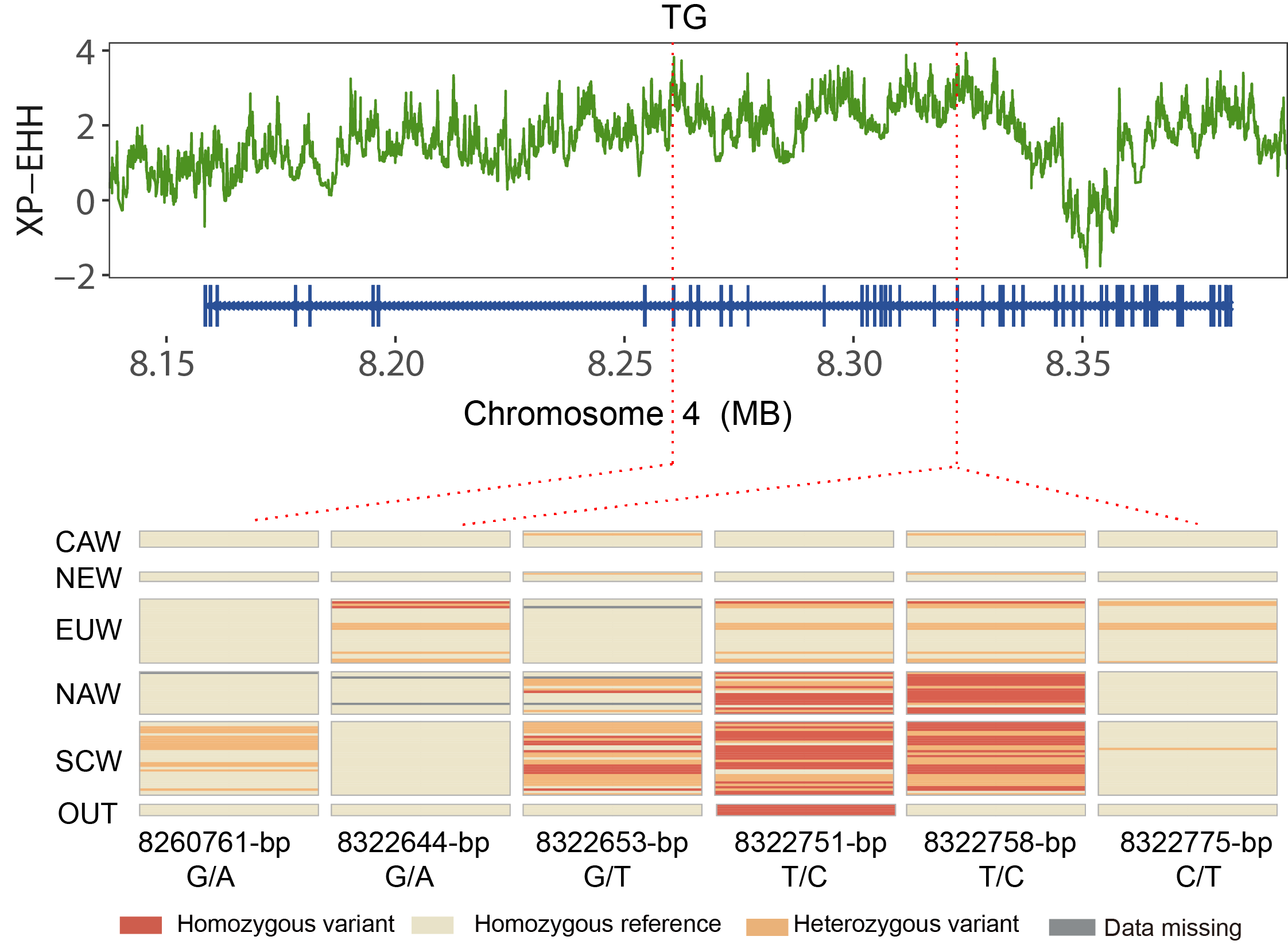

**Figure S17 XP-EHH values around the *TG* gene locus and haplotype pattern of the six missense mutations within the *TG* gene among the OUT and 89 wild boars. OUT: *Sus cebifrons* which were regarded as out group; SCW: Southern Chinese wild boars; NAW: Northeast Asian wild boars; CAW: Central Asian wild boars; EUW: European wild boars; NEW: Near Eastern wild boars.**

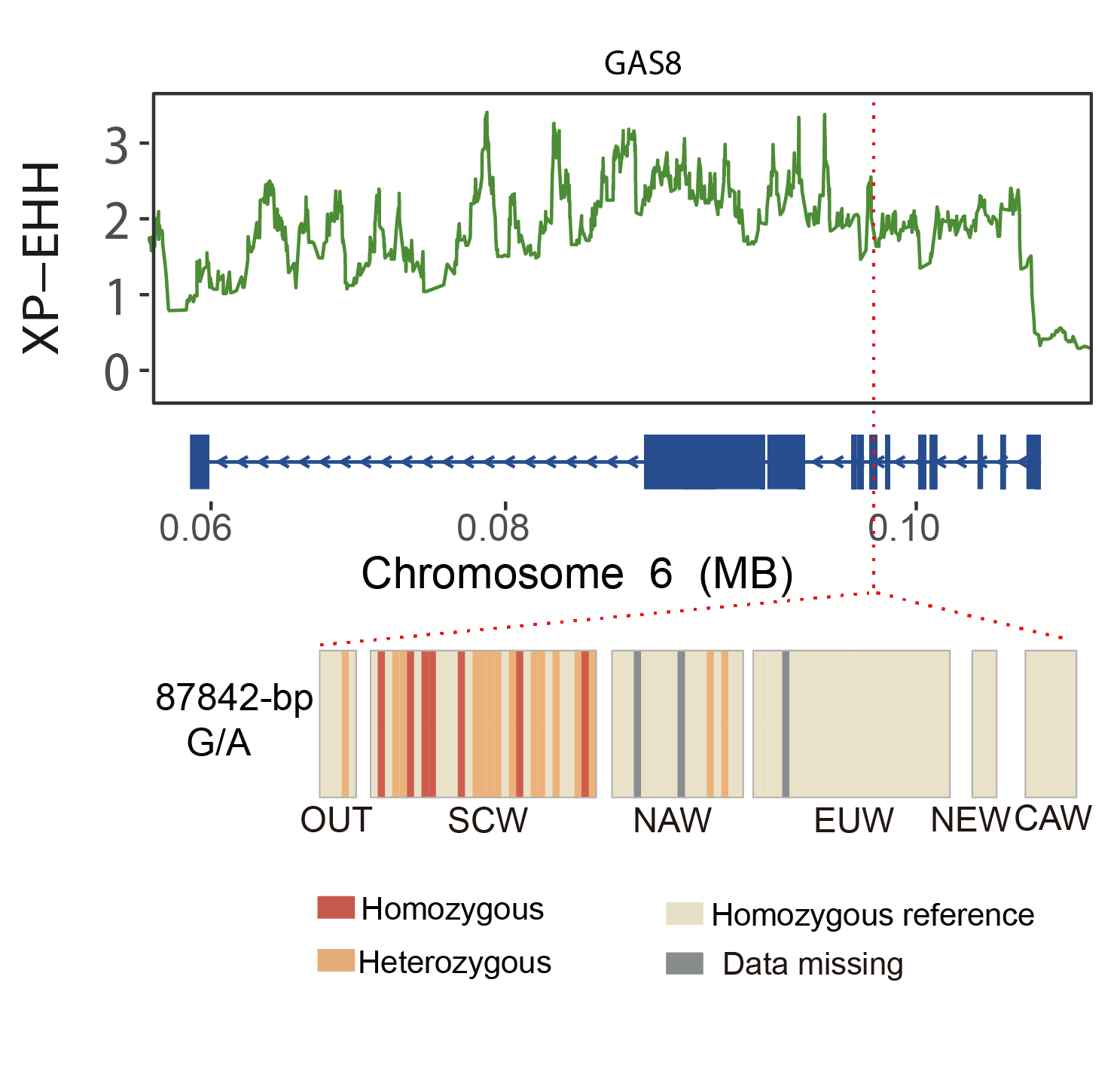

**Figure S18 XP-EHH values around the *GAS8* gene locus and haplotype pattern of the one missense mutation within the *GAS8* gene among the OUT and 89 wild boars. OUT: *Sus cebifrons* which were regarded as out group; SCW: Southern Chinese wild boars; NAW: Northeast Asian wild boars; CAW: Central Asian wild boars; EUW: European wild boars; NEW: Near Eastern wild boars.**

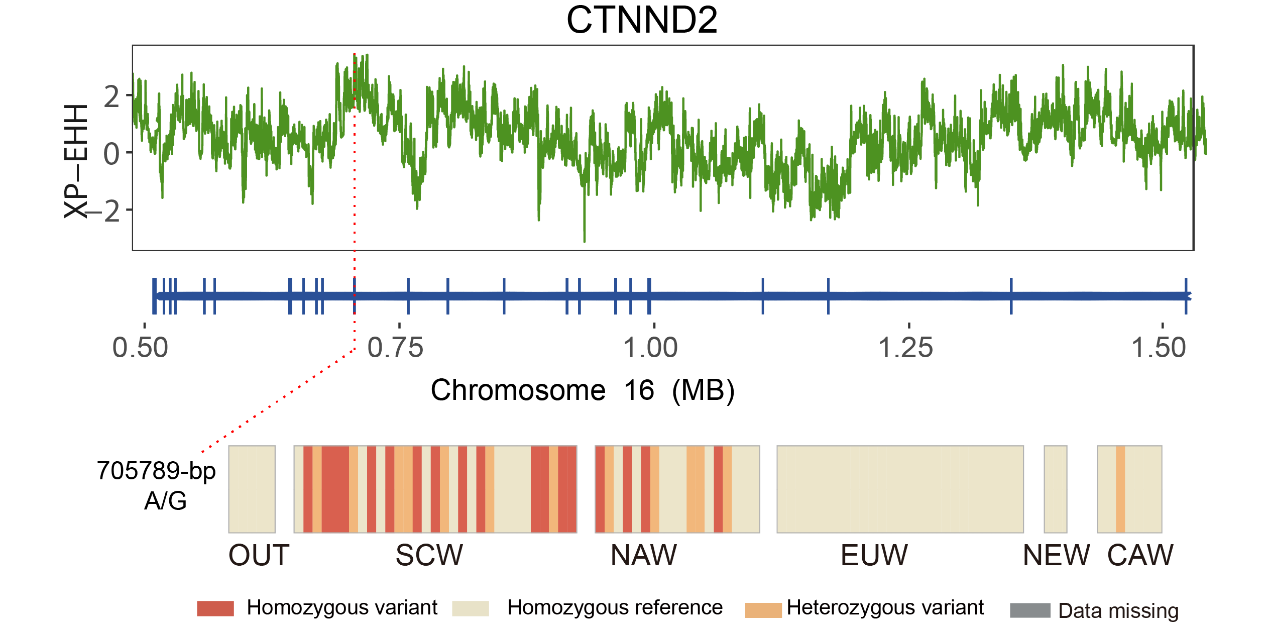

**Figure S19 XP-EHH values around the *CTNND2* gene locus and haplotype pattern of the one missense mutation within the *CTNND2* gene among the OUT and 89 wild boars. OUT: *Sus cebifrons* which were regarded as out group; SCW: Southern Chinese wild boars; NAW: Northeast Asian wild boars; CAW: Central Asian wild boars; EUW: European wild boars; NEW: Near Eastern wild boars.**

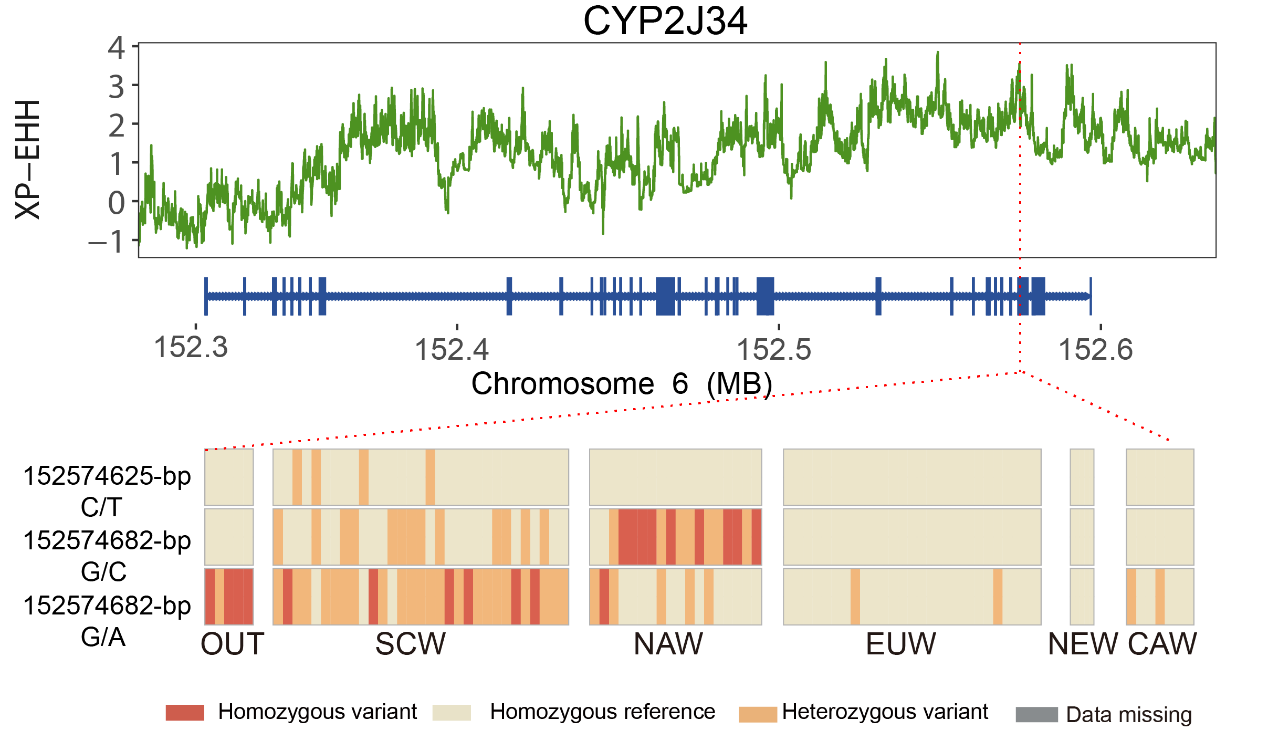

**Figure S20 XP-EHH values around the gene locus and haplotype pattern of the three missense mutations within the *CYP2J34* gene among the OUT and 89 wild boars. OUT: *Sus cebifrons* which were regarded as out group; SCW: Southern Chinese wild boars; NAW: Northeast Asian wild boars; CAW: Central Asian wild boars; EUW: European wild boars; NEW: Near Eastern wild boars.**

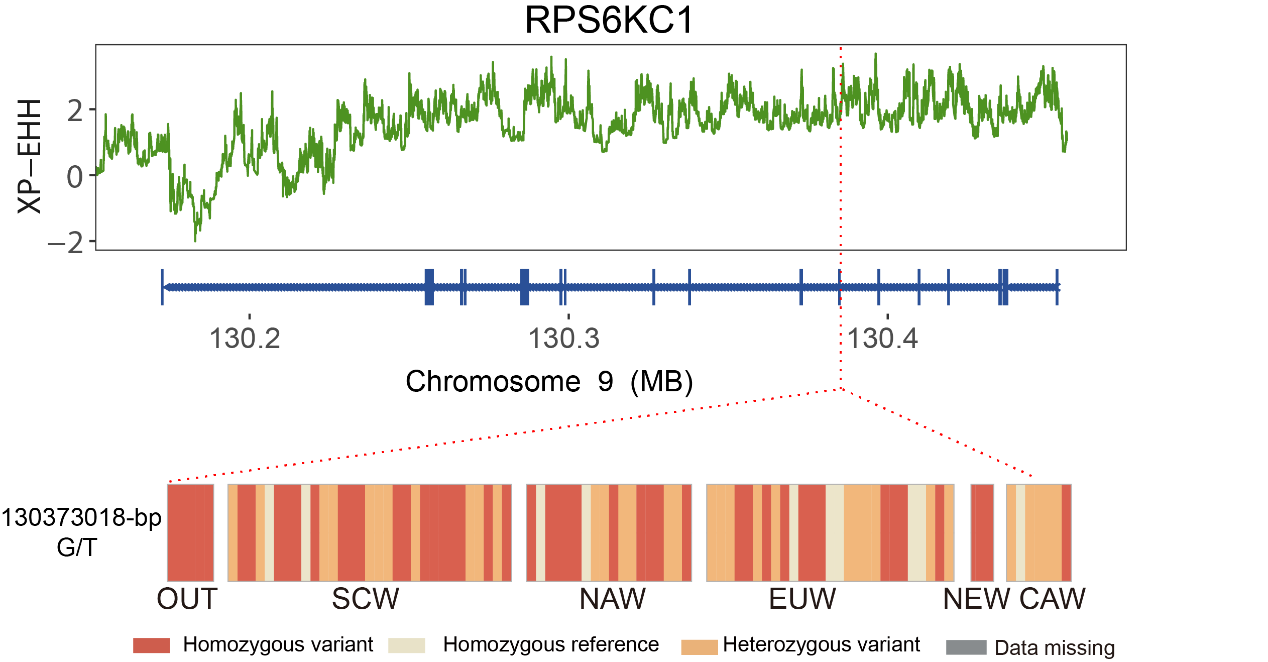

**Figure S21 XP-EHH values around the gene locus and haplotype pattern of the one missense mutation within the *RPS6KC1* gene among the OUT and 89 wild boars. OUT: *Sus cebifrons* which were regarded as out group; SCW: Southern Chinese wild boars; NAW: Northeast Asian wild boars; CAW: Central Asian wild boars; EUW: European wild boars; NEW: Near Eastern wild boars.**

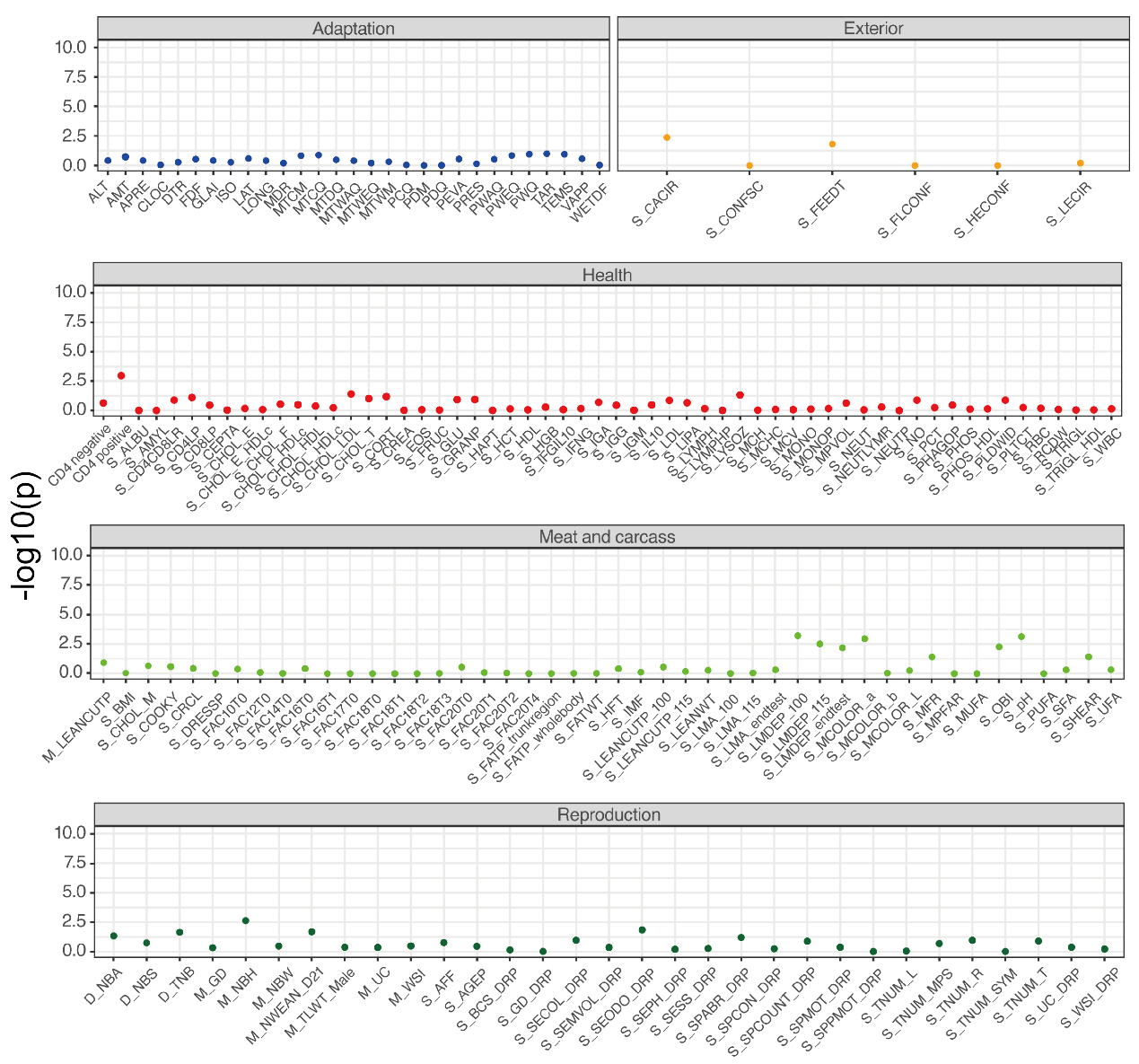

**Figure S22 Point plot shows the association between ALPK2 and complex analysis. Data cite from PigBiobank^2^, each point represents the association between ALPK2 and each trait in gene-based association analysis. The correspondence between the abbreviation and the full name of each trait is in the supplementary table 12.**

1.	Pan Z*, et al.* Pig genome functional annotation enhances the biological interpretation of complex traits and human disease. *Nature Communications* **12**, 5848 (2021).

2.	Zeng H*, et al.* PigBiobank: a valuable resource for understanding genetic and biological mechanisms of diverse complex traits in pigs. *Nucleic acids research* **52**, D980-D989 (2024).
